# Targeted degradation of pathologic tau aggregates via AUTOTAC ameliorates tauopathy

**DOI:** 10.1101/2025.09.10.675324

**Authors:** Jihoon Lee, Su Bin Kim, Hye Yeon Kim, Gee Eun Lee, Dong Won Lee, Eun Hye Jung, Ji Su Lee, Eun Hye Cho, Hong-Beom Park, Da-ha Park, Soon Chul Kwon, Jeong Eun Noh, Hyo Sun Cha, Hans Jin-young Oh, Seung Hoon Lee, Nataša Šterbenc, Cheol Yong Choi, Min Jae Lee, Hyung Min Chi, Young Ho Suh, Kea Joo Lee, Do Hyun Han, Ki Woon Sung, Darja Pavlin, Sung Tae Kim, Chang Hoon Ji, Maja Zakošek Pipan, Yong Tae Kwon

**Affiliations:** Cellular Degradation Biology Center and Department of Biomedical Sciences, College of Medicine, Seoul National University, Seoul 03080, Republic of Korea; AUTOTAC Bio Inc., 225 Gasan digital 1-ro, Seoul, Republic of Korea; Department of Transdisciplinary Medicine, Seoul National University Hospital, 71 Daehakro, Seoul, Republic of Korea; Neuroscience Research Institute, Medical Research Center, Seoul National University, Seoul 03080, Republic of Korea; Neural Circuits Group, Korea Brain Research Institute (KBRI), Daegu 41062, Republic of Korea; Department of Biological Sciences, Sungkyunkwan University, Suwon 16419, Republic of Korea; SGC-UNC, University of North Carolina at Chapel Hill, North Carolina 27599, United States; Department of Chemistry, Pohang University of Science and Technology, Pohang 37673, Republic of Korea; Clinic for Reproduction and Large Animals, Veterinary Faculty, University of Ljubljana, Gerbičeva 60, 1000 Ljubljana, Slovenia; Ischemic/Hypoxic Disease Institute, Medical Research Center, Seoul National University, Seoul 03080, Republic of Korea; Convergence Dementia Research Center, Seoul National University Medical Research Center, Seoul 03080, Republic of Korea; Duke Center for Neurodegeneration Research, Department of Pharmacology and Cancer Biology, Duke Universiry, Durham, United States; Clinic for Small Animals, Veterinary Faculty, University of Ljubljana, Gerbičeva 60, 1000 Ljubljana, Slovenia; Thermo Fisher Scientific, Suseo Office Building 281, Seoul 06349, Republic of Korea

## Abstract

The pathogenesis of tauopathies including Alzheimer’s disease (AD) and progressive supranuclear palsy (PSP) involves the misfolding and aggregation of tau. Here, we employed AUTOTAC to induce the lysosomal degradation of intraneuronal tau aggregates. ATB2005A is a 734-Da chimera that simultaneously binds β-sheet-rich tau aggregates and the autophagic receptor p62/SQSTM1, leading to autophagosomal sequestration and lysosomal co-degradation. In mouse models of tauopathies, orally administered ATB2005A lowered intraneuronal tau aggregates and exerted the therapeutic efficacy in neuroinflammation as well as cognition, behavior, and muscle movements. A Phase 2 clinical trial (U34401-4/2023/14) with companion dogs carrying canine cognitive dysfunction (CCD) demonstrated the efficacy of ATB2005A, as a veterinary medicine, to reverse the disease progression. ATB2005A is under Phase 1 clinical trial with human participants in Korea (202300697). These results validate AUTOTAC as a versatile platform for developing therapeutics to eradicate toxic protein aggregates in a wide range of proteinopathies.

## Main

Tauopathies are driven by misfolded tau aggregates in the brain, leading to the formation of neurofibrillary tangles (NFTs), a hallmark of various diseases, including AD, PSP, and frontotemporal dementia (FTD)^1, 2^. Amongst these, AD is a secondary tauopathy with both tau and amyloid-β (Aβ) deposition and poses a significant health threat with a global prevalence of more than 40 million with 7 million annual cases^3^. AD initially affects the hippocampus and entorhinal cortex, leading to progressive memory loss and spreads to other cortical regions, which is driven by extracellular Aβ plaques and intraneuronal tau deposits^4, 5^. During its pathogenesis, extracellular Aβ initiates a pathophysiological cascade, which drives the hyperphosphorylation of intracellular tau and its misfolding, leading to the formation of NFTs comprising of insoluble tau aggregates^6^. In contrast to AD, PSP is primarily caused by tau aggregation and more acute with the median survival of 5-7 years^7^. PSP is characterized by progressive motor and non-motor symptoms, primarily affecting balance, eye movements, and cognition^7, 8^.

Tau exists in six isoforms, differing in the number of N-terminal inserts and the presence of either three (3R) or four (4R) microtubule-binding repeats^9^. While AD features both isoforms, 4R predominates in PSP and CBD, and 3R in Pick disease (PiD). Despite these differences, all isoforms under pathological conditions share the propensity to be misfolded and form β-sheet-rich oligomers that include heterogeneous species ranging from dimers to larger protofibrillar structures, comprising tens of monomers^10^. These misfolded tau oligomers progressively grow into fibrillar aggregates, also called NFTs^11–13^. NFTs comprise an array of intermolecular β-sheets running parallel to the long axis of the fibrils^12–14^. Compared with insoluble fibrils, smaller, soluble tau oligomers are more neurotoxic^15–17^. Given the pathogenic synergy between Aβ and tau, anti-Aβ therapeutics can reduce early tau alterations but have limited efficacy in late-stage tau pathology^5, 18^.

While traditional approaches to develop therapeutics for AD focused on Aβ, recent studies suggest that tau pathology is more directly responsible for neurotoxicity, neuronal death, and cognitive decline, as the persistent toxicity from existing tau aggregates cannot be simply reversed by reducing Aβ levels^19–21^. For example, the intravenous (IV) administration of donanemab and lecanemab, amyloid-targeting monoclonal antibodies (mAbs), over 1.5 years delayed the disease progression of AD by approximately 35% and 27%, respectively^22–24^. However, constant progression of disease severity persisted along with adverse effects including cerebral edema and microhemorrhages in 24-31% of patients^22–24^. As an alternative approach, targeting pathological tau species sheds light on future directions as the level of NFTs directly correlates to the severity of AD, PSP, and various other tauopathies^25, 26^. Previous studies were mainly undertaken to reduce intracellular tau levels by modulating gene expression or post-translational modifications or extracellular tau by immunotherapy^27–29^. As an effort to eliminate intraneuronal tau, PROTAC (proteolysis-targeting chimera)^30^ has been employed to induce cereblon-mediated ubiquitination and degradation of both phosphorylated and normal tau species in cultured cells^31^. Another tau-PROTAC targeting both wild-type and phosphorylated tau exhibited degradation efficacy primarily to total soluble tau in wild-type mice by intracerebroventricular injection^32^. These degraders focused on tau monomers, limiting their effects on preventing tau aggregation in early stages. Moreover, tau aggregates not only impair proteasomal activity but also cannot go through the inner diameter of the proteasome as narrow as 13 Å, acting as physical limits in exploiting proteasomal degradation in advanced tauopathies^33^.

The N-degron pathway plays a central role in quality control of proteins and cellular constituents through the ubiquitin-proteasome system (UPS) and autophagy-lysosome pathway (ALP)^34, 35^. In the UPS, single N-terminal residues of substrate proteins function as degradation determinants, called N-degrons, which are recognized by cognate N-recognins such as UBR-box containing E3 ligases that induce substrate ubiquitination for proteasomal degradation^36, 37^. We previously identified p62 as an autophagic N-recognin that binds N-degrons via the ZZ domain^38, 39^. This allosteric conformational activation facilitates self-polymerization of p62 in complex with associated cargoes, forming liquid droplets through liquid-liquid phase separation (LLPS). In parallel, the LIR (LC3-interacting region) domain of p62 is exposed to accelerate the delivery of p62-cargo complexes to LC3 on autophagic membranes. Based on the biochemical principle of N-degrons, we developed their chemical mimicries^39, 40^, termed autophagy-targeting ligands (ATLs), that were shown to induce macroautophagic degradation of protein aggregates, cellular organelles, as well as invading pathogens^41^. More recently, we employed the ATL technology to develop AUTOTAC (<u>auto</u>phagy-<u>ta</u>rgeting <u>c</u>himera) that enables targeted degradation of intracellular proteins via p62-dependent macroautophagy^40^. AUTOTAC employs a bifunctional molecule composed of a target-binding ligand (TBL) conjugated to an ATL. A plethora of TBLs can be utilized to enable selective degradation of conventionally undruggable protein targets. As a proof-of-concept approach, a set of Autotacs have been shown to induce lysosomal degradation of disease-associated proteins such as the V7 variant of androgen receptor in castration-resistant prostate cancer and the pathogenic species of α-synuclein in Parkinson’s disease (PD)^42, 43^.

Here, we employed AUTOTAC to develop an oral drug that penetrates the blood-brain barrier (BBB) and induces lysosomal degradation of pathogenic tau species in tauopathies. ATB2005A adopts a derivative of anle138b as a TBL to recognize the β-sheet-rich structures of misfolded tau oligomers and aggregates. In AD and PSP mouse models, oral administration of ATB2005A lowered the levels of insoluble tau species in the brain and exerted the therapeutic efficacy in locomotive activities, behaviors, cognition, and neuromuscular coordination. ATB2005A also reverted the disease progression of dog patients carrying CCD. Our results highlight the therapeutic potential of AUTOTAC to develop disease-modifying drugs for protein aggregates in neurodegenerative diseases.

## Results

### Design of Autotac compounds to degrade pathogenic tau species

To design Autotacs targeting tau aggregates but not its functional monomers, we screened for a TBL that recognizes the β-sheet structure shared by pathogenic tau species in their oligomeric and fibrillar states. A set of compounds, including anle138b, riluzol, ursodeoxycholic acid, 4-phenylbutyrate, resveratrol, baicalein, methylene blue, and curcumin were subjected to thioflavin T (ThT) assays, in which ThT gives fluorescence upon binding to preformed fibrils (PFFs) of aggregation-prone human P301S tau. Anle138b exhibited not only satisfactory pharmacokinetic (PK) profiles and physicochemical properties^44, 45^ but also engagement with PFFs (Supplementary Fig. 1a, b). Consistently, previous studies showed that Anle138b binds the β-sheet structure of α-synuclein oligomers with the dissociation constant (K_d_) of 190 nM^46^. To validate its interaction with PFFs, biotinylated-anle138b associated with tau species was pulled down using streptavidin-coated beads. Non-reducing SDS-PAGE confirmed that biotinylated-anle138b captured increased levels of high molecular weight tau and their dissociated species (Supplementary Fig. 1c). To enhance its druggability and target engagement, anle138b was modified through structure activity relationship (SAR) approach, which resulted in debromination from its para-bromophenyl moiety (Fig. 1a). In ThT assays, debromo-anle138b more effectively engaged with tau fibrils and prevented their self-aggregation. (Supplementary Fig. 1d). To investigate the structural characteristics of this interaction, we employed AlphaFold3 to model the docking of debromo-anle138b onto the structure of tau filaments from AD patient brains (PDB: 8Q8R)^47, 48^. Site-mapping analyses predicted that debromo-anle138b interacts with key residues, including Leu315, Ser316, and Lys317 of tau filaments (Supplementary Fig. 1e-g). The MM/GBSA ΔG binding score for debromo-anle138b was -31.6 kcal/mol, which was more favorable than that of previously known ligands such as X6R (-20.4 kcal/mol) and EGCG (-25.34 kcal/mol)

**Figure 1.**
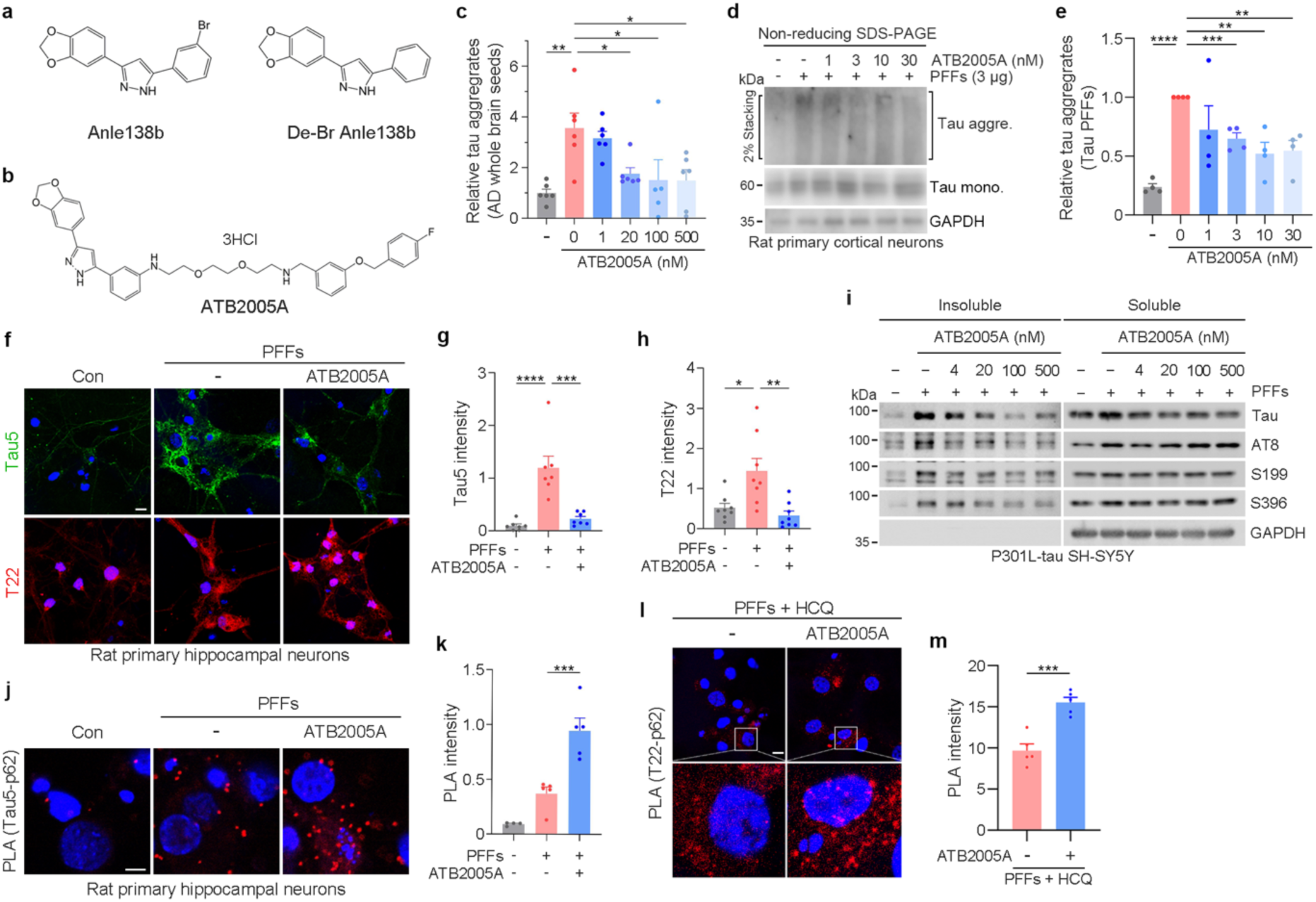
ATB2005A enables targeted degradation of tau aggregates. **(a)** Structures of anle138b and debrominated-anle138b (de-Br-anle138b) compounds. **(b)** Structure of ATB2005A compound. **(c)** ELISA analysis of tau aggregates in HEK293T tau P301S cells treated with 3 μg AD whole brain seeds, followed by treatment with ATB2005A (*n* = 5-6). **(d)** Immunoblotting analysis in rat primary cortical neurons transduced with 3 μg human P301S tau PFFs for 1 week, followed by ATB2005A treatment for 48 h. **(e)** Quantification of (d) (*n* = 4). **(f)** Immunocytochemistry analyses in rat primary hippocampal neurons transduced with 3 μg human P301S tau PFFs for 1 week, subsequently treated with ATB2005A (1 µM, 72 h). The scale bar represents 20 µm. **(g)** and **(h)** Quantifications of (f); *n* = 7 for (g), *n* = 8 for (h). **(i)** Immunoblotting analysis of P301L tau-expressing SH-SY5Y cells following Triton X-100 fractionation assay. 2 μg human P301S tau PFFs were transduced for 24 h, followed by ATB2005A treatment for 24 h. **(j)** Proximity ligation assay in rat primary hippocampal neurons transduced with 3 μg human tau P301S PFFs for 1 week, subsequently treated with ATB2005A (1 µM, 24 h). **(k)** Quantification of (j) (*n =* 5). **(l)** Proximity ligation assay in rat primary hippocampal neurons transduced with 3 µg tau P301S PFF for 1 week, subsequently co-treated with HCQ (10 μM, 24 h) and ATB2005A (1 µM, 24 h). **(m)** Quantification of (l) (*n* = 4). Data are mean ± s.e.m. where relevant. *P*-values (from two-sided unpaired *t* tests): * *P* ≤ 0.05, ** *P <* 0.01, *** *P* < 0.001, **** *P* < 0.0001. Scale bars represent 10 µm otherwise described.

To develop tau-Autotacs, anle138b-based TBL derivatives were conjugated with polyethylene glycol (PEG) linkers in varying topologies, which in turn were conjugated with different ATLs. Following initial screening, an Autotac library was narrowed down to six (ATB2005, ATB2047, ATB2055, ATB2056, ATB2082, and ATB2088), which were further compared with their oral bioavailability and degradation efficacy (Supplementary Fig. 1h; data not shown). We selected ATB2005 with a size of 625 Da from a library of AUTOTACs, which was designed to target protein aggregates in a proof-of-concept study and pathogenic α-synuclein species in PD^40, 43, 51^. To increase its solubility, ATB2005 was formulated into a salt form containing hydrochloride to generate ATB2005A with a size of 734 Da (Fig. 1b). ATB2005A showed improved target engagement with PFFs and physicochemical properties as compared with ATB2005 (Supplementary Fig. 1i, see below).

### ATB2005A enables targeted degradation of tau aggregates

To assess the degradation efficacy of ATB2005A, HEK293T cells expressing human P301S tau, a 4R-type substrate commonly underlying AD and PSP, were treated with brain lysates from AD patients. Homogenous time resolved fluorescence (HTRF) analyses revealed that tau aggregates were degraded by ATB2005A treatment with a half degradation concentration (DC_50_) of ∼20 nM (Fig. 1c). Similar degradation was observed with frontal lobe lysates from AD brains, without observable transcriptional changes of tau (Supplementary Fig. 2a, b). When rat primary cortical neurons were transduced with human P301S tau PFFs (Supplementary Fig. 2c), ATB2005A exhibited DC_50_ of ∼10 nM for lithium dodecyl sulfate (LDS)-resistant tau aggregates (Fig. 1e, f). P301S-tau HEK293T cells transduced with human P301S tau PFFs also exhibited effective degradation of tau aggregates upon treatment with ATB2005A, but not with ATB2005 (Supplementary Fig. 1e, f). Consistently, immunocytochemistry revealed that ATB2005A treatment almost completely eradicated total as well as oligomeric tau species, as visualized with tau5 and T22 antibodies, respectively (Fig. 1f-h). By contrast, ATB2005A lost its degradative activity when autophagy flux was blocked using hydroxychloroquine (HCQ) that inhibits lysosomal acidification (Supplementary Fig. 2d). In sum, ATB2005A induces the degradation of misfolded tau oligomers and their aggregates via the ALP.

To determine the selectivity of ATB2005A for insoluble tau species, SH-SY5Y cells expressing 4R-type P301L tau were transduced with human P301S tau PFFs and fractionated using 1% Triton X-100. Insoluble tau species were markedly eliminated by ATB2005A as determined by immunoblotting analyses using AT8 antibody to phosphorylated-tau (p-tau) at S202 and T205 (Fig. 1i). A similar reduction of insoluble tau species was reproduced using antibodies that recognize S199, T212/S214, or S396 p-tau (Fig. 1i) as well as K18 antibody that recognizes microtubule-binding repeats of tau (Fig. 1i). Despite the reductions of various pathological tau species, no significant changes were observed with soluble tau (Fig. 1i). Thus, ATB2005A induces the degradation of insoluble tau aggregates. Furthermore, immunocytochemistry in P301L-tau SH-SY5Y cells transduced with the PFFs showed that ATB2005A treatment led to recovery of the microtubule meshwork that was otherwise reduced by the PFFs (Supplementary Fig. 2g).

Next, to test whether ATB2005A mediates the engagement of p62 with tau species, we performed *in situ* proximity ligation assay (PLA) in rat primary hippocampal neurons transduced with human P301S tau PFFs. PLA using tau5 and p62 antibodies revealed the recruitment of p62 to tau species as an autophagic quality control (Fig. 1j, k). The basal level of p62-tau engagement was increased by 3-fold upon ATB2005A treatment, which was further enhanced when autophagic flux was blocked using HCQ (Fig. 1j-m). These results demonstrate the selective engagement of ATB2005A to p62.

To assess off-target degradation of cellular proteins by ATB2005A, tandem mass-tag (TMT) proteomics was performed in SH-SY5Y cells expressing human P301L tau following treatment with ATB2005A at 50, 250, and 1000 nM. Out of ∼15,000 proteins capable of being identified, no significant off-target degradation was observed (Supplementary Fig. 2h). As expected, the levels of 50-100 proteins were upregulated, the majority of which were related with the pathways in p62-dependent macroautophagy (Supplementary Fig. 2h). These results suggest that off-target degradation by ATB2005A is at best marginal, given the technical difficulty to recognize a 3D-signature of misfolded tau species lacking normal protein folding.

### ATB2005A is an oral drug that eliminates pathological tau species from the brains of tauopathy model mice

In mice, oral administration of ATB2005A at 10 mg/kg (mpk) exhibited a systemic exposure of 844 ng*h/mL (AUC_last_) with bioavailability of 21%, a half-life (T_1/2_) of 5.3 hr, and brain/plasma ratio of up to 10% at 5 mpk (Supplementary Table 1)^43^. In investigational new drug (IND)-enabling Good Laboratory Practice (GLP) studies (Supplementary Table 2), the no observed adverse effect level (NOAEL) values were at least 240 and 500 mpk in rats and beagle dogs, respectively, upon oral administration 3 times per week (TIW) for 4 weeks. The safety of ATB2005A was also confirmed through examinations on respiration, central nervous system, DNA integrity, and cardiac functions (Supplementary Table 2). ATB2005A is under Phase 1 clinical trials at Seoul National University Hospital in South Korea with 76 healthy volunteers (202300697).

To evaluate the efficacy of ATB2005A in AD and PSP mouse models, we used JNPL3 mice expressing human P301L tau in the brains^52^. These mice exhibit deficits in behavior, cognition, and neuromuscular functions from 5 months of age, providing a model for progressive behavioral decline^53, 54^. The mice at the age of 7 months were intraperitoneally (IP) injected with 10 mpk ATB2005A 5 times per week for 6 weeks. Immunohistochemistry of the brain sections using AT8 antibody specific to S202/T205 p-tau revealed the formation of NFTs in ectorhinal, perihinal, and motor cortices, which were all markedly lowered by ATB2005A (Fig. 2a, b). The degradation of detergent-insoluble tau aggregates was reproduced using immunoblotting of whole brain lysates (Fig. 2c, d). Thus, ATB2005A induces the degradation of pre-existing NFTs from the brains of mice.

**Figure 2.**
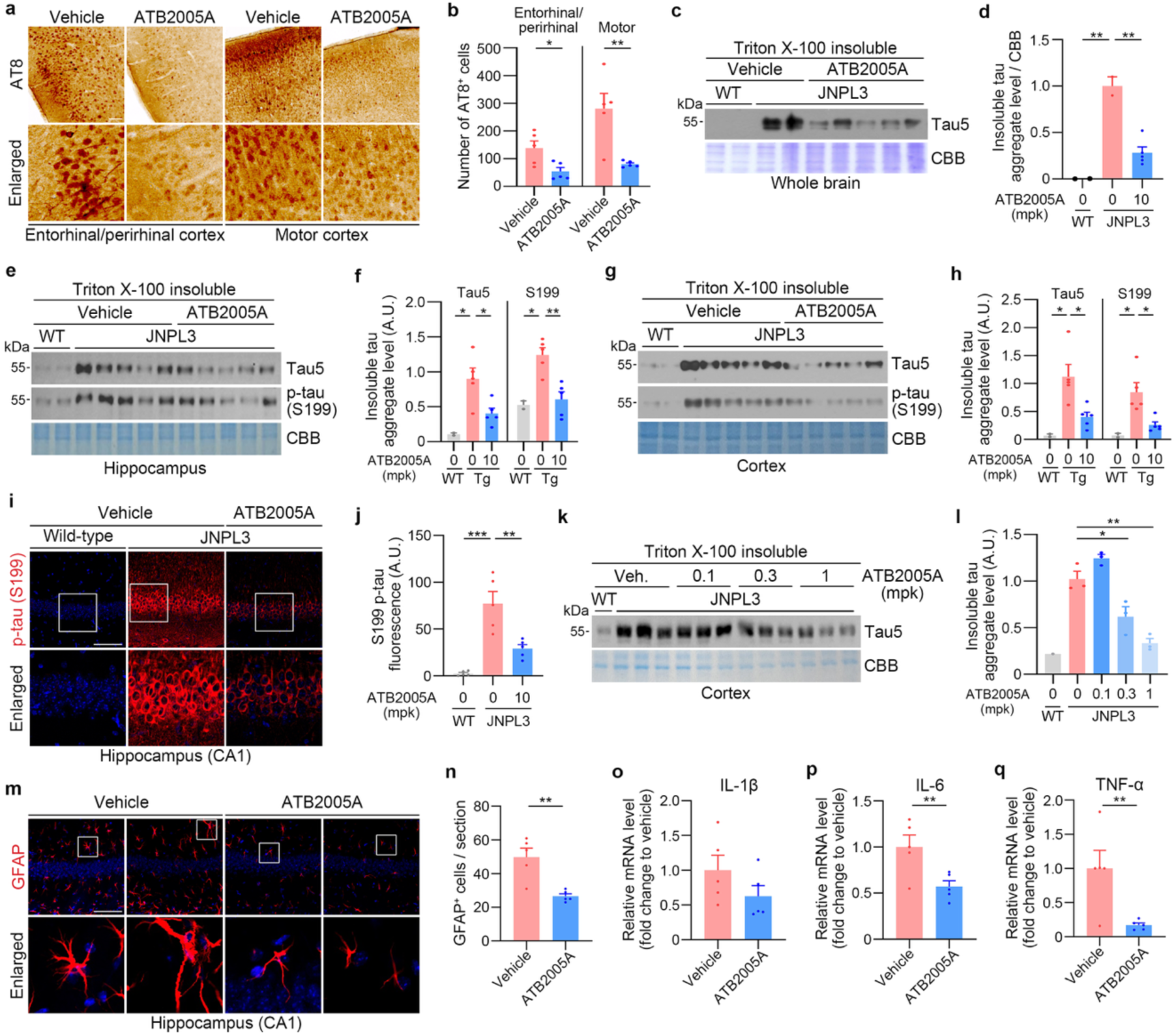
ATB2005A is an oral drug that eliminates pathological tau species from the brains of tauopathy model mice. **(a)** Immunohistochemistry analysis of cortical regions in JNPL3 mice intraperitoneally injected with vehicle or 10 mpk ATB2005A. **(b)** Quantification of AT8^+^ cells in (a) (*n* = 5). **(c)** Immunoblotting analysis of Triton X-100 insoluble fractions of the wild-type and JNPL3 mouse brains. The mice were intraperitoneally injected with vehicle or 10 mpk ATB2005A **(d)** Quantification of insoluble tau aggregate levels in (c) (*n* = 2 for wild-type and vehicle*, n* = 5 for ATB2005A). **(e)** and **(g)** Immunoblotting analyses of Triton X-100 insoluble fractions of the hippocampus and the cortex. The mice were orally administered with vehicle or 10 mpk ATB2005A. **(f)** and **(h)** Quantifications of insoluble tau aggregate levels in (e) and (g) (*n* = 2 for wild-type, *n* = 5 for vehicle and ATB2005A). **(i)** Immunofluorescence analysis of S199 p-tau in the hippocampus of wild-type and JNPL3 mice. The mice were orally administered with vehicle or 10 mpk ATB2005A. **(j)** Quantification of S199 p-tau^+^ fluorescence levels in (i) (*n =* 5). **(k)** Immunoblotting analysis of Triton X-100 insoluble fractions of wild-type and JNPL3 mouse cortices. The mice were orally administered with vehicle or 0.1, 0.3, and 1 mpk ATB2005A. **(l)** Quantification of insoluble tau aggregate levels in (k) (*n =* 1 for wild-type, *n* = 3 for each dose). **(m)** Immunofluorescence analysis of GFAP^+^ cells in the hippocampus of JNPL3 mice intraperitoneally administered with vehicle or 10 mpk ATB2005A. **(n)** Quantification of (m) (n = 5). **(o**–**q)** Relative mRNA levels of IL-ϕ3, IL-6, and TNF-α in the whole brains of JNPL3 mice, intraperitoneally administered with vehicle or 10 mpk ATB2005A. Data are mean ± s.e.m. where relevant. *P*-values (from two-sided unpaired *t* tests): * *P* ≤ 0.05, ** *P <* 0.01, *** *P* < 0.001. Scale bars represent 100 µm.

To develop an oral tau-AUTOTAC, 10-month-old JNPL3 mice were orally administered by 10 mpk ATB2005A TIW for 8 weeks. Immunoblotting analyses of cortex and hippocampal proteins showed that aggregated species of total tau and S199 p-tau were markedly reduced by ATB2005A administration (Fig. 2e-h), without significant changes in soluble tau species in the cortex (Supplementary Fig. 3a, b). Immunofluorescence analyses of hippocampal slices also revealed the lowering of S199 p-tau inclusions induced by ATB2005A (Fig. 2i, j). We also assessed dose-dependence in 12-month-old JNPL3 mice orally administered with ATB2005A at 0.1, 0.3, and 1 mpk TIW for 8 weeks. ATB2005A induced the degradation of insoluble total tau species at as low as 0.3 mpk and more significantly at 1 mpk (Fig. 2k, l). Thus, ATB2005A is an oral drug that targets tau aggregates but not their functional monomers from tauopathy model mice.

### ATB2005A exerts therapeutic efficacy in the behavior, cognition, and neuromuscular functions in tauopathy model mice

To determine the therapeutic efficacy of ATB2005A in cognitive and behavioral deficits, JNPL3 mice were orally administered with 5 mpk ATB2005A TIW for 8 weeks. In open field tests, ATB2005A-treated mice exhibited 95% increase in the total distance, an indicator of locomotive activity and exploratory behavior, as compared with the vehicle group (Fig. 3a, b). Robust improvements were further supported by the distance (15-fold) and time (20-fold) in the center zone that measure anxiety-associated behavior (Fig. 3c, d). In 6-arm radial maze tests that monitor spatial learning and memory, ATB2005A administration alleviated the cognition declines as determined by latency to first correct choice (13.3 vs. 5.6 sec) and trial to second correct choice (4.7 vs 1.8 sec) (Fig. 3e-g). These results demonstrate the efficacy of ATB2005A in learning, memory, and anxiety-associated behavior.

**Figure 3.**
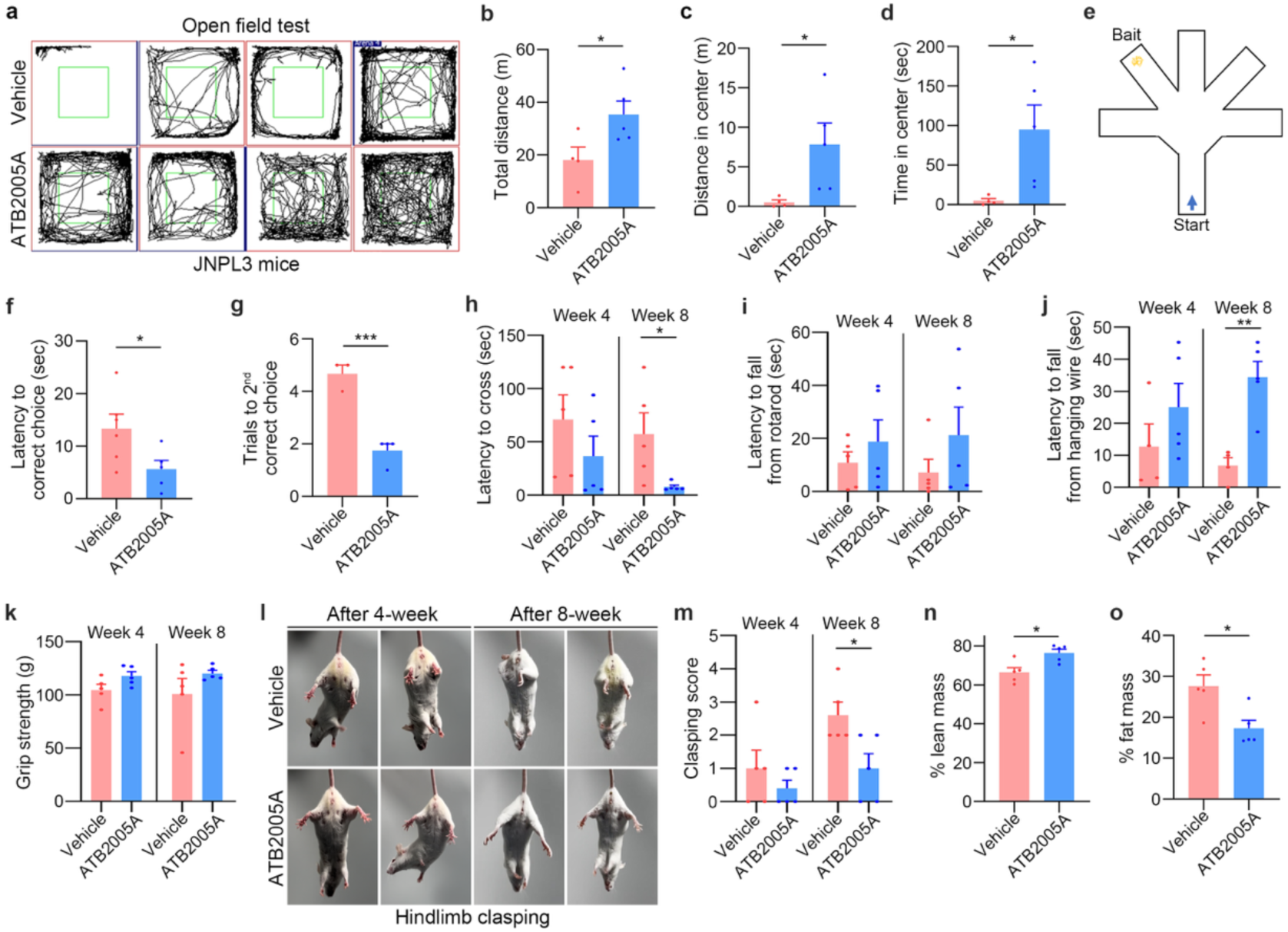
ATB2005A exerts therapeutic efficacy in the behavior, cognition, and neuromuscular functions in tauopathy model mice. **(a)** Open field test in JNPL3 mice orally administered with vehicle or 5 mpk ATB2005A (TIW for 8 weeks). **(b**–**d)** Quantifications of total distance traveled, distance in center, and time in center in (a) (*n* = 4–5). **(e)** Schematic of the 6-arm radial maze. **(f)** and **(g)** Quantifications of the latency to first correct choice and the number of trials to second correct choice in (e); *n* = 5–6 for (f), *n* = 3–4 for (g). **(h)** Latency to cross in the beam walking test after 4- or 8-week treatment of ATB2005A (*n* = 5). **(i)** Latency to fall from the rotarod test (*n* = 5). **(j)** Latency to fall in the hanging wire test (*n* = 5). **(k)** Forelimb muscle strength in the grip strength test (*n* = 5). **(l)** Representative images of hindlimb clasping behavior. **(m)** Quantification of clasping score in (l); *n* = 5. **(n)** and **(o)** Body composition analyses by EchoMRI showing percentages of lean mass and fat mass (*n* = 5). Data are mean ± s.e.m. where relevant. *P*-values (from two-sided unpaired *t* tests): * *P* ≤ 0.05, ** *P <* 0.01, *** *P* < 0.001.

Tauopathy progressively aggravates neuromuscular deficits as prominently manifested in PSP, FTD, and CBD^2, 9^. To assess the efficacy of ATB2005A in movement dysfunctions, we conducted behavioral examinations on JNPL3 mice orally administered with 10 mpk ATB2005A TIW for 8 weeks. ATB2005A-treated mice exhibited 85% reduction of the latency to cross in beam-walking tests (Fig. 3h) and 3-fold reduction of the time to fall in rotarod tests (Fig. 3i, Supplementary Video 1). These improvements in balance and motor coordination correlated to 5-fold reduction of the latency to fall in hanging wire tests that measure grip strength, muscle endurance, and neuromuscular coordination (Fig. 3j). Consistently, grip strength tests showed 16% increase in forelimb muscle strength (Fig. 3k). It is known that JNPL3 mice with motor deficits tend to clasp their hindlimbs when suspended instead extending them outwards^55^, which was reduced by 62% by ATB2005A (Fig. 3l, m). JNPL3 mice also develop dystonia, a form of abnormal muscle contraction, also known as gait ignition failure (GIF). Notably, 15 oral doses of 10 mpk ATB2005A prevented the muscle dysfunction and the onset of GIF (see Supplementary Video 2). These findings demonstrate that ATB2005A not only prevents neuromuscular decline but also actively preserves and enhances motor function in tauopathy model mice.

### ATB2005A mitigates neuroinflammation and improves body content in tauopathy model mice

The accumulation and propagation of tau aggregates underlie glial hyperactivation and neuroinflammation in AD brains with tau pathology^56^. To determine the efficacy of ATB2005A in astrocyte hypertrophy, 6-month-old JNPL3 mice were IP injected with 10 mpk ATB2005A 5 times per week for 6 weeks. Immunofluorescence analyses of the brain sections revealed the accumulation of GFAP^+^ hypertrophic astrocytes and insoluble tau aggregates in the hippocampus, both of which were markedly reduced by ATB2005A (Fig. 2m, n). We also assessed its effects on systemic neuroinflammation in the brains of JNPL3 mice. RT-qPCR analyses of the brains revealed that ATB2005A significantly reduced the levels of pro-inflammatory cytokines, including IL-1β, IL-6, and TNF-α (Fig. 2o-q). These results demonstrate that the removal of neurotoxic tau species alleviates neuroinflammation in tauopathies.

Tauopathies are often accompanied by systemic metabolic alterations leading to changes in body weight and fat distribution^57^. To determine the impact of tau degradation on body compositions, we measured percentages of fat and lean mass by whole body scanning of the JNPL3 mice. EchoMRI analyses showed that as compared with vehicle treatment, ATB2005A treatment reduced fat content by 38% and instead increased the percentage of lean mass by 15%, indicative of the gain of muscle mass (Fig. 3n, o). These suggest that the lowering of NFTs may improve body composition in tauopathies.

### ATB2005A induces targeted degradation of tau aggregates in a mouse model of severe early-onset tauopathy

To determine the efficacy of ATB2005A to mitigate severe tauopathy, we adopted rTg4510 mice that express human P301L tau at 13 times the level of endogenous murine tau^58^. This accelerated model of advanced tauopathy exhibits the deficits in cognition and spatial memory from 3-4 months and neuronal loss by 5 months^59, 60^. Male mice at the age of 4 months orally received 1 or 10 mpk ATB2005A TIW for 4 months, which did not affect body weights (Fig. 4a, Supplementary Fig. 4a). Immunoblotting of brain RIPA lysates revealed the accumulation of total as well as S199, S202/T205, S212/T214, and S396 p-tau species in the cortex and hippocampus (Fig. 4b, c, Supplementary Fig. 4b, c), which were all significantly lowered by ATB2005A treatment at 1 mpk and to a greater extent at 10 mpk. The degradative efficacy was also reproduced at both doses using immunostaining of S202/T205 p-tau (Fig. 4d, e, Supplementary Fig. 4d, e). The selectivity of ATB2005A to tau aggregates was confirmed in immunoblotting of sarkosyl-insoluble proteins of the cortex and hippocampus using antibodies to total tau and S202/T205 p-tau (Fig. 4f-i, Supplementary Fig. 4f-i). These results further validate the degradative efficacy of ATB2005A for insoluble tau species in tauopathies.

**Figure 4.**
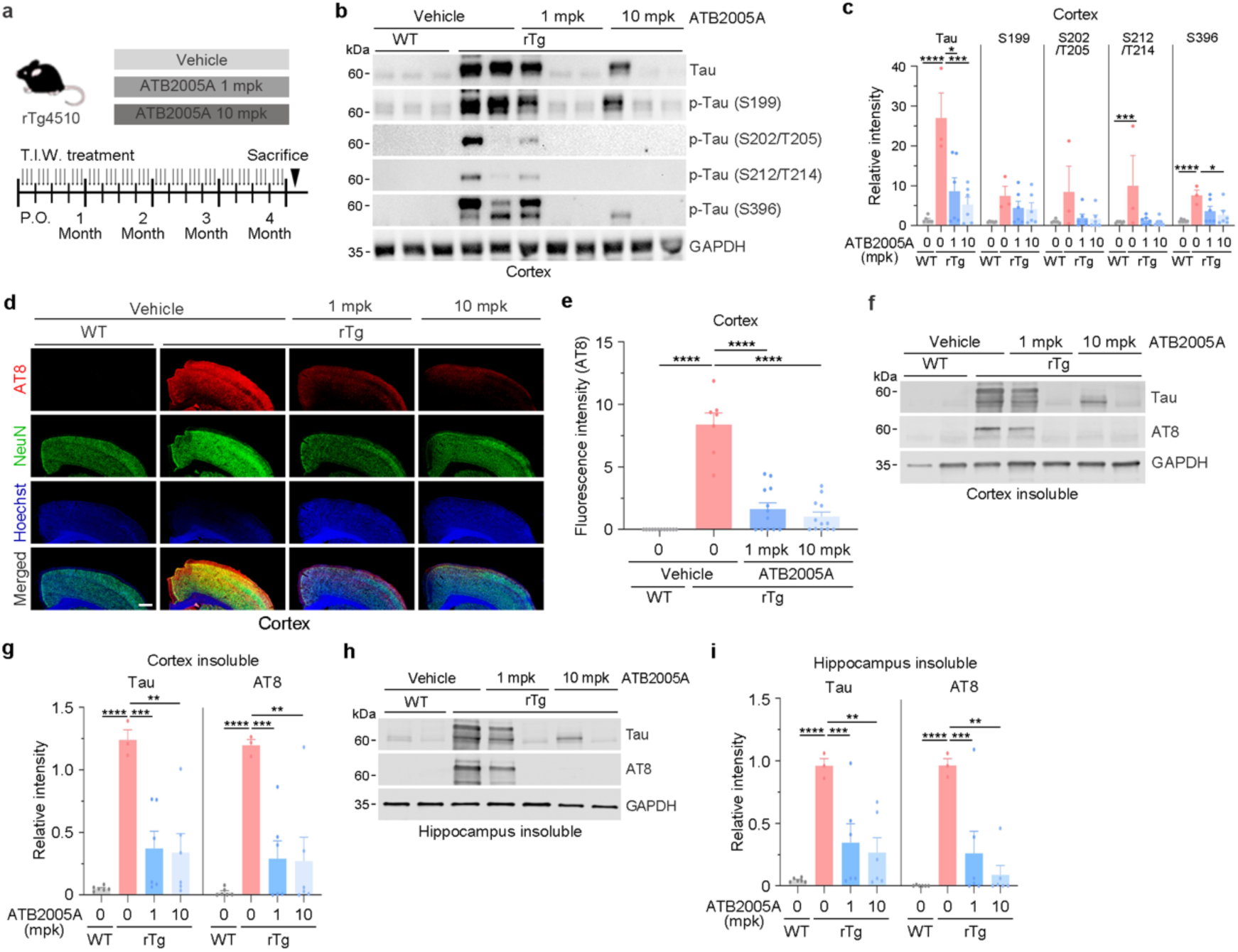
ATB2005A induces targeted degradation of tau aggregates in a mouse model of severe early-onset tauopathy. **(a)** Oral administration schedule and doses of ATB2005A in the rTg4510 mouse model. These mice were orally treated with 1 or 10 mpk ATB2005A three times per week for 4 months. **(b)** Immunoblotting analyses of the cortical lysates of wild-type and rTg4510 mice. Various pathologic p-tau species were monitored. **(c)** Quantification of total tau and p-tau (S199, S202/T205, S212/T214, and S396) in (b); *n =* 3 or 6. **(d)** Immunofluorescence analyses of S202/T205^+^ p-tau in the cortex of wild-type and rTg4510 mice. The scale bar represents 200 µm. **(e)** Quantification of (d) (*n =* 7 or 12). **(f)** and **(h)** Immunoblotting analyses of sarkosyl-insoluble cortical or hippocampal lysates of wild-type and rTg4510 mice. Insoluble total tau and S202/T205^+^ p-tau aggregates were monitored. **(g)** and **(i)** Quantifications of (f) and (h), respectively; *n =* 3 or 6. Data are mean ± s.e.m. where relevant. *P*-values (from two-sided unpaired *t* tests): * *P* ≤ 0.05, ** *P <* 0.01, *** *P* < 0.001, **** *P* < 0.0001.

### ATB2005A ameliorates behavioral deficits in rTg4510 mice

To examine whether ATB2005A treatment rescues rTg4510 mice from locomotor hyperactivity during disease progression associated with tauopathy^59^, the mice orally administered at 1 or 10 mpk ATB2005A TIW for 4 months were subjected to open-field tests (Fig. 4a). Vehicle-treated rTg4510 mice developed tauopathy-associated hyperactivity as indicated by 2.3-fold increase in traveling distances as compared with wild-type mice (Fig. 5a, b). By sharp contrast, mice treated with both doses exhibited almost normal behavioral patterns, including travelling distances, demonstrating the efficacy of ATB2005A in locomotor hyperactivity. To monitor spatial learning memory, we performed Morris water maze tests. ATB2005A treatment improved the reference memory at 1 mpk and more significantly at 10 mpk, the latter of which improved 60% and 100% in the time spent in target quadrants and the number of entries, respectively (Fig. 5c-f). This correlated to 90% increase in the number of platform crossings with 80% decrease in the latency to reach the platform (Fig. 5g, h). When the learning curve was assessed throughout 6 days, ATB2005A treatment improved spatial learning by 50% (Fig. 5i), demonstrating that ATB2005A ameliorates spatial memory and learning deficits in the severe tauopathy. Finally, we determined whether ATB2005A improves recognition and working memory using novel object recognition tests, in which ATB2005A increased the discrimination index by 2-fold to the level similar to that of wild-type mice (Fig. 5j, k, and Supplementary Fig. 4j). These results further validate the therapeutic efficacy of ATB2005A to reverse deficits of recognition memory in tauopathy.

**Figure 5.**
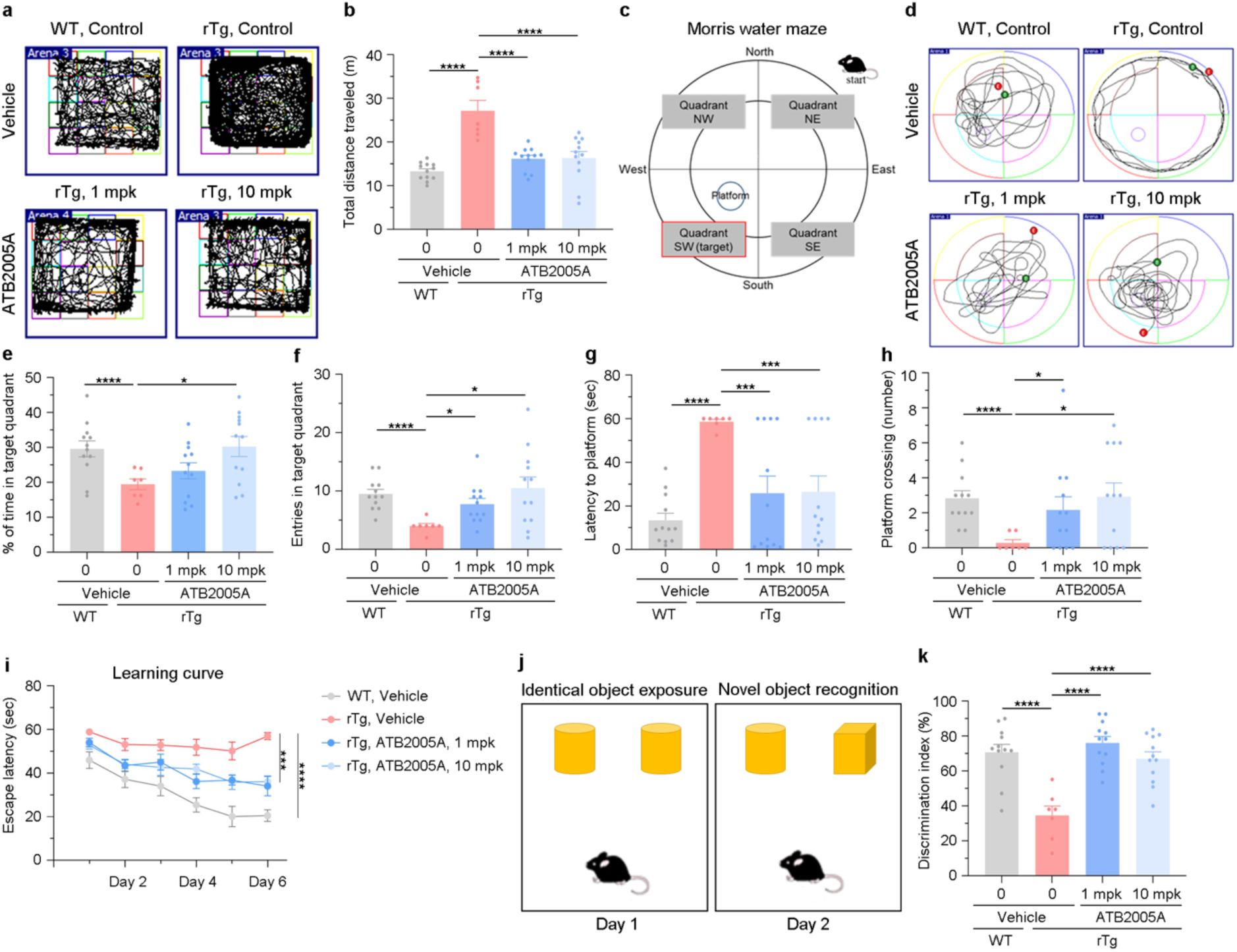
ATB2005A ameliorates behavioral deficits in rTg4510 mice. **(a)** Open field test in wild-type and rTg4510 mice. These mice were orally treated with 1 or 10 mpk ATB2005A three times per week for 4 months. **(b)** Quantification of total distance traveled in (a); *n* = 7– 12 mice per group. **(c)** Schematic diagram of the Morris water maze test. **(d)** Morris water maze test in wild-type and rTg4510 mice orally administered with vehicle or ATB2005A (1 mpk or 10 mpk). **(e**–**h)** Quantifications of time in target quadrant, entries into target quadrant, latency to platform, and platform crossing in (d); *n* = 7–12 mice per group. **(i)** Learning curve of wild-type and rTg4510 mice orally administered vehicle or ATB2005A (1 mpk or 10 mpk); *n* = 7– 12 mice per group. **(j)** Schematic diagram of the novel object recognition test. **(k)** Quantification of discrimination index in (j); *n* = 7–12 mice per group. Data are mean ± s.e.m. where relevant. *P*-values (from two-sided unpaired *t* tests): * *P* ≤ 0.05, *** *P* < 0.001, **** *P* < 0.0001.

### ATB2005A exhibits therapeutic efficacy to ameliorate CCD in a Phase 2 clinical trial with companion dogs

The deposition of hyperphosphorylated tau aggregates is a direct cause of CCD, which is accompanied by behavioral deficits in social interactions, sleep-wake cycles, learning, and memory^61, 62^. To validate the effectiveness and safety of ATB2005A as a veterinary drug to treat CCD, we conducted a double-blind, randomized, placebo-controlled, Phase 2 clinical trial following ethical approval (Approval No. U34401-4/2023/14) from the Ethical Committee of the Ministry of Agriculture, Forestry and Food, Veterinary Administration of the Republic of Slovenia (UVHVVR) on May 16^th^, 2023, for implementation at the University of Ljubljana’s Veterinary Faculty. Dogs were eligible for inclusion if they were within the last 25% of their expected lifespan according to the AKC (American Kennel Club) breed standards and displayed CCD symptoms based on behavioral examinations (see Methods). Total 32 companion dogs were enrolled and equally divided into drug and placebo groups, the former orally receiving 15 mpk ATB2005A capsules TIW for 6 weeks (Fig. 6a, b, Supplementary Table 3). No adverse effects of ATB2005A were observed in complete blood count and biochemical examinations for serum concentrations of urea, creatine, total proteins, albumins, total bilirubin, and the serum activities of alanine aminotransferase, alkaline phosphatase, aspartate transaminase, and glutamyl transferase (Supplementary Table 4).

**Figure 6.**
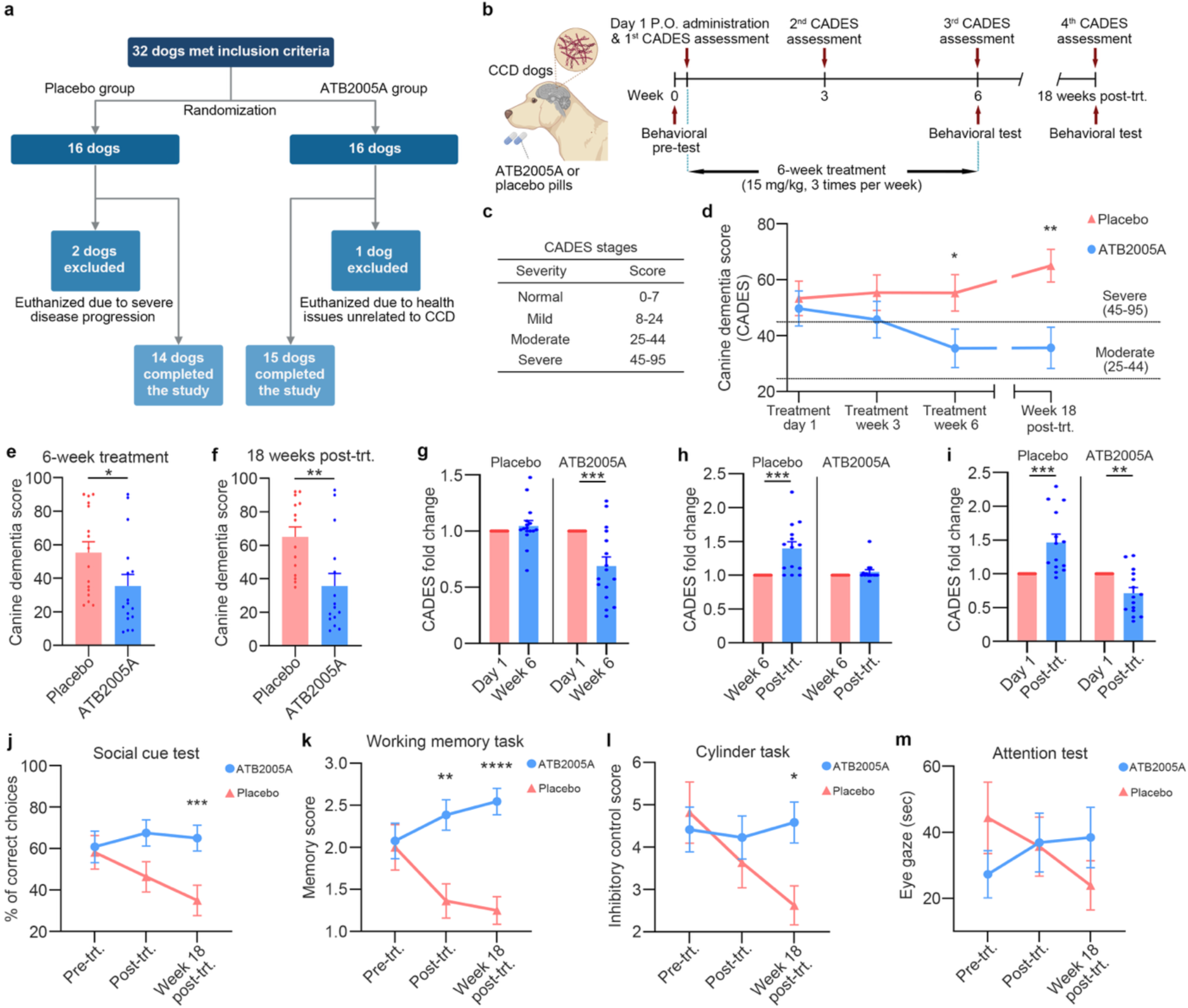
ATB2005A exhibits therapeutic efficacy to ameliorate CCD in a Phase 2 clinical trial with companion dogs. **(a)** Flowchart showing the study design. A total of 32 companion dogs with CCD were randomized into placebo (*n* = 16) and ATB2005A (*n* = 16) treatment groups; 14 and 15 dogs completed the study, respectively. **(b)** Schematic of the experimental timeline. Dogs received oral administration of ATB2005A (15 mpk, three times per week) or placebo for 6 weeks. CADES and behavioral tests were assessed at baseline (week 0), during treatment (week 3 and 6), and at the follow-up time point (week 18 post-treatment). **(c)** CADES ranges to define dementia severity. **(d)** CADES at different time points of ATB2005A or placebo group. **(e)** CADES after the 6-week treatment (*n* = 16 each). **(f)** CADES after 18 weeks post-treatment (*n* = 14 for placebo, *n* = 15 for ATB2005A). **(g)** Fold-changes of CADES at week 6 from day 1 (*n* = 16 each). **(h)** Fold-changes of CADES at week 18 post-treatment from the completion of treatment (*n* = 14–16). **(i)** Fold-changes of CADES scores at 18 weeks post-treatment from day 1 (*n* = 14–16). **(j–m)** Behavioral tests showing enhanced social recognition (*n* = 10–12), working memory (*n* = 11–13), inhibitory control (*n* = 8–12), and sustained attention (*n* = 10–13) in ATB2005A-treated dogs. Data are mean ± s.e.m. where relevant. *P*-values (from two-sided unpaired *t* tests): * *P* ≤ 0.05, ** *P <* 0.01, *** *P* < 0.001, **** *P* < 0.0001.

To assess changes in cognitive functions and disease severity, the CADES (canine dementia scale) was monitored based on owner assessments that comprise behavioral changes in spatial orientation, social interactions, sleep-wake cycles, and house soiling^62^. On day 1, the mean CADES values for the placebo and ATB2005A groups were 53 and 50, respectively, both at severe dementia stages (Fig. 6c, d). When the fold-change of CADES was assessed at week 6 from day 1, the ATB2005A group significantly improved the dementia score (*P* = 0.0006), whereas the placebo group worsened. This demonstrates that 18 doses of ATB2005A reversed the disease severity, to a moderate stage within the CADES range of 25 to 44 (Fig. 6d, e). Consistently, a significant improvement of CADES was observed when the disease trajectory of each group was directly compared at week 6 (*P* = 0.0445) (Fig. 6g). Next, when the follow-up assessment was conducted at week 18 post-treatment, the CADES of the placebo group was worsened from the end of the administration period (*P* = 0.0002) (Fig. 6f, h, i). Notably, ATB2005A-treated dogs showed no significant decline during the post-administration period. In addition, a direct comparison of post-treatment trajectories between the two groups revealed a significant improvement (*P* = 0.0049) (Fig. 6f). These results demonstrate that ATB2005A exhibits therapeutic efficacy to reverse severe CCD and sustain prolonged cognition improvement.

To evaluate the efficacy of ATB2005A for behavioral and cognitive deficits in CCD, we performed a series of cognitive tests for social cues, working memory, cylinder task, and sustained attention. When the recognition memory was determined through the social cue test, ATB2005A treatment for 6 weeks increased the number of correct choices as compared to the placebo group (Fig. 6j). This efficacy was sustained for 18 weeks following the completion of treatment, as indicated by a significant improvement (*P* = 0.0062). Next, the working memory task was tested by hiding treats with containers for progressive delays. ATB2005A treatment significantly improved the memory score when compared to the placebo group at week 6 (*P* = 0.0011, which was maintained during the post-treatment period as indicated by a more robust enhancement (*P* < 0.0001) (Fig. 6k). Next, the cylinder test was performed to assess the task memory and inhibitory control by requiring dogs to retrieve treats from a transparent cylinder. ATB2005A treatment improved the inhibitory control score at week 6 (*P* > 0.05) and significantly at week 18 post-treatment (*P* = 0.0123) (Fig. 6l), that otherwise constantly decreased by over 6 months in the placebo group. In attention test that measures the duration of eye contact to assess sustained attention, ATB2005A treatment improved the mean duration of eye gaze from 24 to 37 sec, which was sustained during the post-treatment period (Fig. 6m). However, a consistent decline in sustained attention was observed in the placebo group. These cumulatively demonstrate the disease-modifying efficacy of ATB2005A in the companion dogs carrying CCD.

## Discussion

The rising prevalence of neurodegenerative diseases poses a growing challenge to global health, particularly as societies enter a super-aging era^63^. The devastating conditions, including but not limited to AD, PSP, PD, and ALS, progressively rob individuals of cognition, motor functions, and most importantly, their independence^64^. Although tauopathies represent the major class of neurodegenerative diseases, no fundamental cure currently exists^18^. Recent advancements have introduced mAbs against Aβ into the market, though their efficacy and safety profiles are controversial. Donanemab slowed cognitive and functional decline by 35% over 76 weeks in early symptomatic AD patients with low to medium tau burden, but was associated with a high incidence (∼24%) of amyloid-related imaging abnormalities (ARIA)^22^. Lecanemab targets soluble Aβ protofibrils and showed a 27% reduction in clinical decline over 18 months, but similarly exhibited ARIA-E (edema) and ARIA-H (hemosiderin deposition) in 13.6% and 16.0% of treated patients^23^. Moreover, the intracellular localization of tau remains a major obstacle to antibody-based therapies, prompting a shift in focus toward tau-targeting strategies. Crucially, mounting evidence suggests that tau, not Aβ, is more directly associated with neurodegeneration and clinical progression in AD^18^. A recent longitudinal PET imaging study demonstrated that while Aβ accumulation may initiate disease, regional tau spreading correlates more closely with symptom onset and disease acceleration^65^. This underscores the therapeutic relevance of lowering tau pathology to slow or halt disease progression, even in the presence of residual amyloid burden.

Here, we employed AUTOTAC to develop ATB2005A, a BBB-penetrating oral drug that enables the lysosomal degradation of pathologic tau aggregates via p62-dependent macroautophagy. This strategy is rooted in the N-degron pathway, a degradative system in which N-degrons are recognized as degradation signals^35^. We developed ATLs that mimic Arg/N-degrons and allosterically activate p62, promoting selective cargo sequestration and autophagosome formation^40^. Conjugation of these ATLs to tau-binding ligands yielded ATB2005A, which selectively targets β-sheet-rich tau aggregates without affecting functional monomers (Fig. 1). In tauopathy mouse models, ATB2005A exerted the degradation activities for insoluble tau aggregates but not its functional monomers as well as the therapeutic efficacy in cognition, memory, neuromuscular coordination, locomotive activities, and muscle strength as well as neuroinflammation and body contents (Fig. 2–Fig. 5). In Phase 2 trial with companion dogs suffering from CCD to evaluate the effectiveness and safety of ATB2005A as a veterinary medicine, ATB2005A not only exhibited therapeutic efficacy in the deficits of behavior, cognition, and memory but also reversed the diseases progression. In GLP toxicity studies, ATB2005A demonstrated its safety as exemplified by NOAEL of 500 mpk in beagle dogs orally administered TIW for 4 weeks. As a human medicine, ATB2005A is under Phase 1 clinical trials. This study is the first to demonstrate the therapeutic approach that eliminates the causative protein aggregates underlying a broad spectrum of neurodegenerative diseases, affecting ∼100 million people.

Together, these findings position AUTOTAC as a mechanistically distinct and clinically promising strategy that directly eliminates tau aggregates and offers a novel therapeutic avenue for a broad spectrum of neurodegenerative diseases.

## Methods

The detailed description of materials and methods used in this study is included in the Supplementary Information.

## Data availability

All data are available from the corresponding authors upon request.

## Acknowledgements

This work was supported by grants from the National Research Foundation (NRF) of Korea funded by the Ministry of Science, ICT, and Future Planning (MSIP) (NRF-2020R1A5A1019023 to Y.T.K., M.J.L., D.H.H., and H.M.C., and NRF-2021R1A2B5B03002614 to Y.T.K.; RS-2021-NR059245 and RS-2024-00332875 to M.J.L.), the Korea Health Industry Development Institute and Korea Dementia Research Center (KDRC) funded by the MSIP (RS-2024-00447844 to C.H.J. and Y.T.K.; RS-2023-00261784 to M.J.L.; RS-2024-00332875 to Y.H.S.), the Korea Brain Research Institute (KBRI) Basic Research Program (25-BR-01-03 to K.J.L), the Slovenian Research Agency (P4-0053 to M.J.P.), and the NRF funded by the Ministry of Education (RS-2023-00249464 to C.H.J.; RS-2024-00446110 to S.B.K.; and RS-2024-00461291 to C.H.J. and S.B.K.; RS-2024-00412741 to H.Y.K.).

## Contributions

Y.T.K. supervised the study. J.L., S.B.K., H.Y.K., E.H.J., E.H.C., J.S.L., S.T.K., and C.H.J. designed, performed, and analyzed *in vitro* experiments; D.W.L, J.L., and K.W.S. designed, performed, and analyze experiments with JNPL3 mice; J.E.N. and K.J.L. designed, performed, and analyzed experiments with rTg4510 mice; D.H.P. and Y.H.S. provided primary rat hippocampal and cortical neurons; H.J.O., S.H.L., and H.M.C. synthesized Autotac compounds and biotinylated TBLs; J.L., G.E.L, N.S., D.P., and M.Z.P. designed, performed, and analyzed experiments with CCD dogs; J.L., S.B.K., and H.Y.K. visualized the data of the manuscript; J.L., S.B.K., H.Y.K., and Y.T.K. wrote original and revised manuscript; H.S.C., C.Y.C., M.J.L., Y.H.S., S.T.K., C.H.J., H.B.P., N.S., D.P., and M.Z.P., and D.H.H proofread and edited the manuscript; All authors discussed the results and commented on the manuscript.

## Corresponding authors

Correspondence to Sung Tae Kim, Chang Hoon Ji, Maja Zakošek Pipan, and Yong Tae Kwon

## Ethics declarations

The authors declare that they have no competing interests.

## Supplementary Figures & Figure legends

**Supplementary Figure 1.**
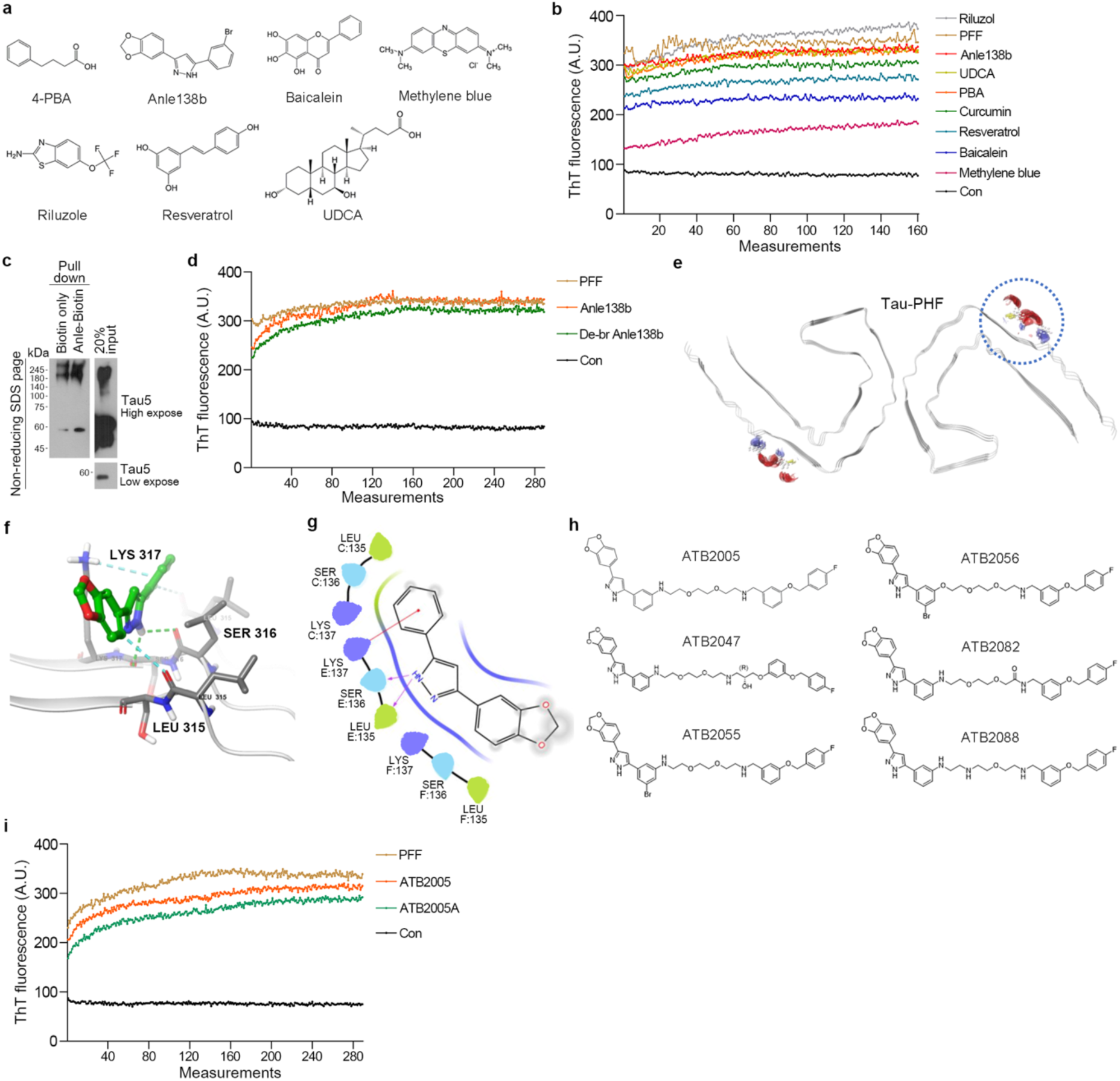
Characteristics of TBLs and Autotac compounds. **(a)** Molecular structures of 4-PBA, anle138b, baicalein, methylene blue, riluzole, resveratrol, and UDCA. **(b)** ThT assay with human P301S tau PFFs and the series of TBLs. 5 μg tau PFFs and 5 μM of compounds were incubated with 2.5 μM ThT solution with constant agitation of 300 rpm for 40 h. Each measurement was taken every 15 min. **(c)** Pull-down assay with 5 μg human P301S tau PFFs and biotinylated anle138b. **(d)** ThT assay with human P301S tau PFFs and anle138b or de-Br anle138b. 5 μg tau PFFs and 1 μM of compounds were incubated with 2.5 μM ThT solution with constant agitation of 300 rpm for 72 h. Each measurement was taken every 15 min. **(e)** AlphaFold3-predicted docking conformation of de-Br anle138b on the cryo-EM structure of tau PHFs from AD patient brains (PDB: 8Q8R). **(f)** Enlarged view of de-Br anle138b-interacting residues Leu315, Ser316, and Lys317. **(g)** Two-dimensional interaction diagram illustrating predicted molecular binding between de-Br anle138b and the tau-PHF binding pocket. **(h)** Molecular structures of ATB2005, ATB2047, ATB2055, ATB2056, ATB2082, and ATB2088. **(i)** ThT assay with human P301S tau PFFs and ATB2005A or ATB2005. 5 μg tau PFFs and 1 μM of the compounds were incubated with 2.5 μM of ThT solution with constant agitation of 300 rpm for 72 h. Each measurement was taken every 15 min.

**Supplementary Figure 2.**
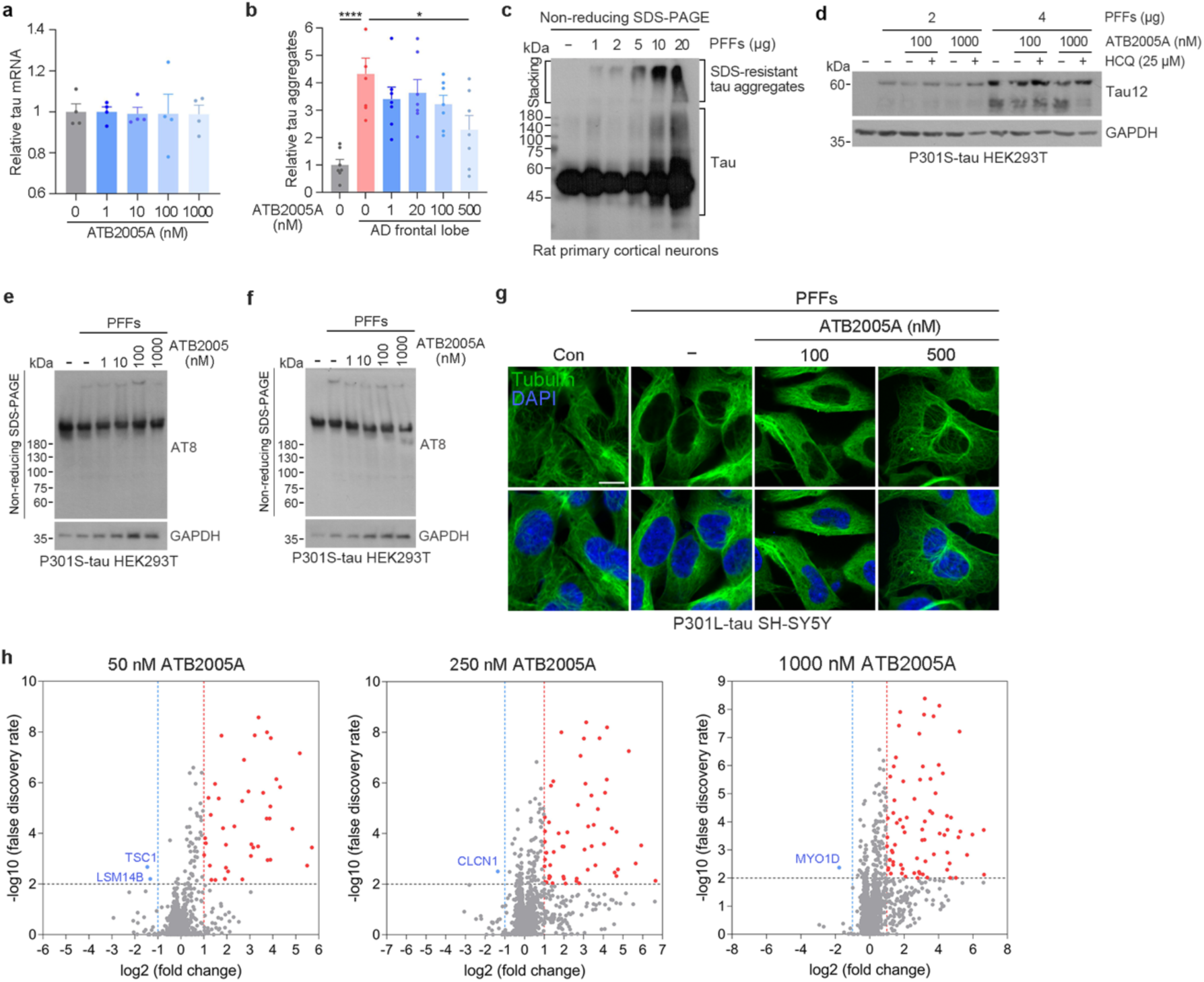
ATB2005A induces degradation of insoluble tau aggregates without significant off-target degradation. **(a)** Relative tau mRNA levels in SH-SY5Y P301L tau cells upon treatment of ATB2005A (24 h) (*n* = 4). **(b)** ELISA analysis of tau aggregates induced by treatment of 3 μg AD frontal lobe seeds into HEK293T P301S tau cells, followed by treatment with ATB2005A for 24 h (*n* = 6–7). **(c)** Oligomerization assay in rat primary cortical neurons transduced with human P301S tau PFFs for 1 week. **(d)** Immunoblotting analysis in HEK293T P301S tau cells transduced with human P301S tau PFFs (24 h), followed by ATB2005A (24 h) and HCQ (24 h) treatment. The tau aggregates were completely dissociated by sonication in RIPA buffer. **(e)** and **(f)** Non-reducing SDS-PAGE in HEK293T P301S tau cells transduced with human P301S tau PFFs (24 h), followed by ATB2005 or ATB2005A treatment (48 h). ATB2005A induced the degradation of high molecular-weight tau aggregates whereas ATB2005 did not. **(g)** Immunocytochemistry in SH-SY5Y P301L tau cells transduced with human P301S tau PFFs (100 nM, 48 h), subsequently treated with ATB2005A (24 h). **(h)** TMT-labelled proteomic LC-MS was performed in SH-SY5Y P301L tau cells treated with ATB2005A at 50, 250, and 1000 nM for 24 h.

**Supplementary Figure 3.**
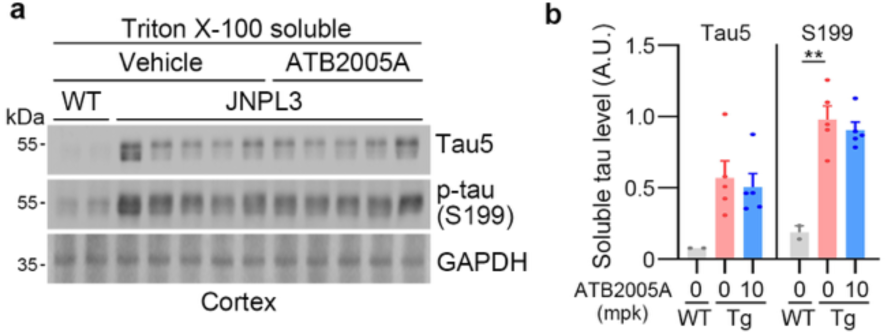
Soluble tau species are not targeted by ATB2005A in mouse brains with tauopathy. **(a)** Triton X-100 soluble fractions in the cortex of wild-type and JNPL3 mice orally administered with vehicle or ATB2005A (10 mpk). **(b)** Quantification of soluble tau levels in (a) (*n =* 2–5).

**Supplementary Figure 4.**
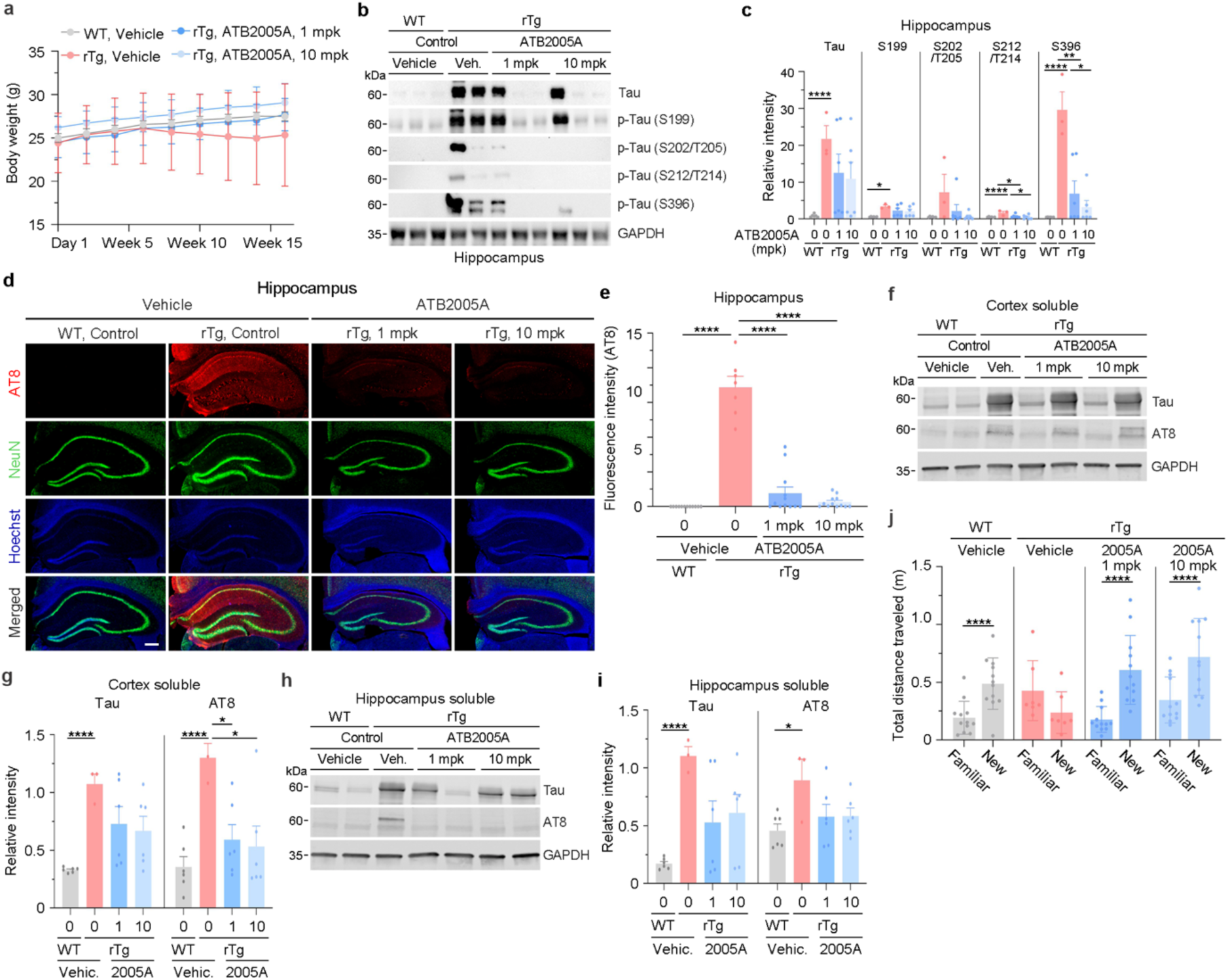
ATB2005A ameliorates behavioral deficits in rTg4510 mice. **(a)** Body weight in wild-type and rTg4510 mice orally administered with vehicle or ATB2005A (1 mpk or 10 mpk) (*n =* 3 or 6). **(b)** Immunoblotting of hippocampal lysates of wild-type and rTg4510 mice orally administered with vehicle or ATB2005A (1 mpk or 10 mpk). **(c)** Quantifications of tau and p-tau (S199, S202/T205, S212/T214, and S396) species levels in (b) (*n =* 3 or 6). **(d)** Immunocytochemistry in the hippocampus of wild-type and rTg4510 mice orally administered with vehicle or ATB2005A (1 mpk or 10 mpk). The scale bar represents 200 µm. **(e)** Quantification of (d) (*n =* 7 or 12). **(f)** and **(h)** Immunoblotting with sarkosyl soluble fractions of cortices and hippocampi in wild-type and rTg4510 mice orally administered with vehicle or ATB2005A (1 mpk or 10 mpk). **(g)** and **(i)** Quantifications of (f) and (h); *n =* 3 or 6. **(j)** Novel object test to measure total distance traveled near the new object for exploration. Wild-type and rTg4510 mice were orally administered with vehicle or ATB2005A (1 mpk or 10 mpk) (*n =* 7 or 12). Data are mean ± s.e.m. where relevant. *P*-values (from two-sided unpaired *t* tests): * *P* ≤ 0.05, ** *P <* 0.01, **** *P* < 0.0001.

