## Supplementary Information for "Targeted degradation of pathologic tau aggregates via AUTOTAC ameliorates tauopathy"

**Methods and materials**

**Canine patient recruitment**

Recruitment efforts included collaboration with veterinary clinics across Slovenia, distribution of informational brochures to veterinarians and dog owners, email campaigns, and announcements on the website of Veterinary Faculty at the University of Ljubljana. Additional outreach involved lectures on canine cognitive dysfunction (CCD) at veterinary events, newspaper advertisements, and word-of-mouth referrals. Dogs were eligible for inclusion if they were within the last 25% of their expected lifespan according to American Kennel Club (AKC) breed standards and displayed CCD symptoms. Additional criteria included no swallowing difficulties, stable chronic conditions for at least six months, a consistent diet for at least two months, and private ownership. Exclusion criteria comprised systemic illness that could interfere with their cognitive function, aggressive dogs, neurological disorders requiring psychoactive medications, blindness, immobility, seizures, systemic or metabolic diseases affecting the central nervous system, chronic liver or kidney disease based on laboratory results, and cardiac arrhythmias. In addition, individuals receiving pharmacological agents or supplements (e.g. selegiline hydrochloride, nicergoline, propentofylline, or dietary regimens) commonly prescribed for cognitive dysfunctions or age-related symptoms were ineligible for inclusion. To standardize baseline conditions, dogs assigned to the experiment were required to discontinue such diets or interventions targeting cognitive decline for a minimum of 30 days prior to enrollment and throughout the study period.

**Clinical trial design**

The experimental protocol complied with the ethical standards laid down in the European Directive 2010/63/EU on animal experimentation. Written informed consent was obtained from all participating owners before the start of the experimental treatment. The clinical study received ethical approval (Approval No. U34401-4/2023/14) from the Ethical Committee of the Ministry of Agriculture, Forestry and Food, Veterinary Administration of the Republic of Slovenia (UVHVVR) on May 16, 2023, for implementation at the University of Ljubljana’s Veterinary Faculty from October 2023 through December 2026. A total of 32 client-owned dog patients that met the prespecified inclusion criteria were enrolled in a double-blind, randomized, placebo-controlled trial. Dogs were randomly assigned to either the placebo group (*n* = 16) or the ATB2005A treatment group (*n* = 16). Treatments were administered orally three times per week for 6 weeks using either ATB2005A pills or visually indistinguishable placebo controls. Both investigators and owners remained blinded to group allocation throughout the trial. Cognitive function was assessed using the Canine Dementia Scale (CADES) at baseline, after 3 weeks and 6 weeks of treatment, and again at 18 weeks post-treatment. Behavioral testing was conducted at baseline, after 6 weeks of treatment, and at the 18-week post-treatment follow-up.

**Clinical evaluations and CADES monitoring**

Owners and veterinarians at the veterinary faculty completed a comprehensive questionnaire covering dog and owner demographics, environmental factors, behavior, diet, medications, and overall health status. Cognitive function was assessed by CADES scoring, comprising behaviors in spatial orientation, social interaction, sleep-wake cycles, and house-soiling. Clinical evaluations included full physical, ophthalmologic, orthopedic, and neurological examinations, along with complete blood cell count determined by an automated laser hematology analyzer ADVIA 120 (Siemens, Munich, Germany). Biochemical examinations on serum concentrations of urea, creatinine, total proteins, albumins, total bilirubin, and the serum activities of alanine aminotransferase (ALT), alkaline phosphatase (AP), aspartate transaminase (AST) and gamma-glutamyl transferase (GGT) were performed with automated biochemistry analyzer RX Daytona (Randox, Crumlin, UK). No adverse effects of ATB2005A were observed in complete blood count and biochemical parameters.

**Cognitive tests**

Cognitive tests were performed according to 2019 AAHA Canine Life Stage Guidelines. Briefly, the social cue tasks began with a warm-up phase where dogs learned to retrieve treats from under cups, followed by pointing and marking trials (10 each) and an odor control test (8 trials). The result was presented as percentage of correct choices. The working memory task involved hiding treats under cups with progressively longer delays (6-120 sec). The number of correct decisions at each delay and the longest delay with at least 4 out of 6 correct decisions was recorded for each dog. Based on these results, dogs were assigned one of three grades; grade 1: <20 sec, grade 2: 20-60 sec, grade 3: >60 sec. The cylinder test (8 trials) assessed inhibitory control by requiring dogs to retrieve treats from a transparent cylinder without touching the sides, followed by a detour task where their preferred side was blocked. Results were expressed as the percentage of missions successfully completed. The sustained attention was measured by recording the duration of eye contact with an experimenter. The duration of sustained eye contact was recorded. The test was repeated three times for each dog, and an average duration was analyzed.

**Thioflavin T fluorescence assay**

Total 5 μg of recombinant tau preformed fibrils (PFFs; Novus Biologics, NBP2-76794) were incubated with test ligands at a designated concentration in reaction buffer (0.01 M HEPES and 0.1M NaCl in distilled water) containing 2.5 μM Thioflavin T (ThT), in a total volume of 100 μL per well. The reactions were conducted in a 96-well plate at 37 °C with continuous agitation at 300 rotations per minute (rpm) in the plate reader. ThT fluorescence (excitation/emission: 440/485 nm) was measured every 15 min for up to 72 h.

**LC-MS/MS sample preparation and analysis**

P301L tau-expressing SH-SY5Y cells were seeded at 5.0 × 10⁶ cells per 100-mm dish and treated with ATB2005A at 50, 250, and 1000 nM for 24 h. Following treatment, cell pellets were lysed with RIPA buffer (Cell Nest, CNR001-0100). Protein concentrations were determined using the BCA protein assay kit (Thermo Fisher Scientific, 23225). For each sample, 200 µg of protein was subjected to filter-aided sample preparation (FASP) digestion. Peptides were desalted using C18 microspin columns (Harvard Apparatus), preconditioned with 100% methanol, 0.1% formic acid, and 80% acetonitrile in 0.1% formic acid. Samples were loaded, washed with 0.1% formic acid (50 µL), eluted with 100 µL of 80% acetonitrile, and vacuum-dried.

For proteomic analysis, 100 µg of peptide was labeled using the TMT 16-plex isobaric label reagent set (Thermo Fisher Scientific) and fractionated using a C18 Sep-Pak column with a gradient of acetonitrile in 0.1% triethylamine. 20 fractions were combined into 10 and vacuum-dried using a HyperVac Max VC2200 (Hanil Scientific). Peptides were separated by nano-LC with a trapping column (75 µm × 2 cm, C18) and an analytical column (75 µm × 50 cm, PrpMap RSLC C18). The mobile phases consisted of solvent A (water with 0.1% formic acid) and solvent B (80% acetonitrile with 0.1% formic acid). A multistep gradient of solvent B was applied at 300 nL/min. MS spectra were acquired over m/z 400–2000. Data were processed using Proteome Discoverer software (version 2.5, Thermo Fisher Scientific) against the UniProt human database. Quantification was performed using TMT reporter ions with modifications: TMTpro (N-terminus and lysine), oxidation (methionine), acetylation (protein N-terminus), carbamidomethylation (cysteine), and Met-loss+acetyl. Statistical analyses were performed using one-way ANOVA (individual protein level) followed by Tukey’s HSD post-hoc test.

**Protein and ligand preparation for docking analysis**

Crystal structures of α-synuclein (PDB ID: 7WMM) and microtubule-associated protein tau (PDB ID: 8Q8R, 7P65) were obtained from Protein Data Bank (PDB). Proteins were processed and refined using protein preparation wizard in Maestro (Schrödinger 14.2)^1^, including the addition of missing hydrogen atoms, correction of bond orders, and energetic optimization using the OPLS_2005 force field. The root mean square deviation (RMSD) of the heavy atom was set to 0.3Ǻ during refinement.

The synthesized compounds Anle138b and ATB2005A were converted from 2D to 3D structures and optimized using LigPrep in Maestro. The process included correction of protonation and ionization states at pH 7.4, removal of salts, addition of hydrogens, and generation of tautomeric and ionization variants. Final structures were energy-minimized using the OPLS_2005 force field with a molecular mechanics energy function and an RMSD cutoff of 0.01 Å to generate the low-energy ligand isomer.

**Sitemap analysis**

Potential ligand-binding sites were identified using the SiteMap tool (v2.6, Schrödinger LLC)^2^. Parameters included a restrictive definition of hydrophobicity, 15 site points per reported site, and a maximum of 15 reported sites. Predicted sites were visually inspected to select those within relevant binding regions. If multiple sites were found in the same region, the site with the highest SiteScore was retained. For each selected site, residues within 3.5 Å of the site points were extracted and saved as a separate PDB file for further analysis.

**Receptor grid generation and molecular docking**

Grid boxes were defined by selecting the co-crystallized ligand (1KI and X6R of 7WMM, and 8UQ7, respectively), ensuring consistent box dimensions centered on each ligand. After ligand removal, binding sites were validated using SiteMap, confirming alignment with the original ligand positions. Docking was performed using Glide in standard precision (SP) mode with the OPLS_2005 force field^3^. A rigid receptor protocol was applied, keeping the protein fixed while allowing ligands flexibility.

**Prime MM/GBSA**

The molecular mechanics with generalized Born surface area (MM-GBSA) method was used to estimate the relative binding free energy (ΔG bind) of protein-ligand and protein-protein complexes, using the Prime^1^ module. Complexes were energy-minimized with the OPLS_2005 force field prior to calculation. Binding free energy was calculated using the VSGB solvation model and the OPLS_2005 force field, based on the following equation:

ΔG = ΔG_solvation + ΔE_minimized
 + ΔG_surface

Ligand-protein interactions were visualized using the ligand interaction diagram tool in Maestro.

**Cell culture**

The Tau RD P301S FRET Biosensor cell line (ATCC, CRL-3275) was purchased from ATCC. The SH-SY5Y cells expressing P301L mutant tau-GFP were purchased from Innoprot. The cells were cultured in Dulbecco’s Modified Eagle’s Medium (Life Technologies, Gibco, 11995-065) supplemented with 10% FBS in a standard 37 °C, 5% CO_2_ humidified incubator.

**Primary neuron culture**

All procedures were approved by the Seoul National University Institutional Animal Care and Use Committee (IACUC) and conducted in accordance with institutional guidelines. Primary cortical and hippocampal neurons were prepared from embryonic day 18 (E18) Sprague Dawley rat embryos. Pregnant rats were anesthetized and euthanized by CO_2_ inhalation, and embryos were collected immediately. Dissections were carried out in dissociation buffer containing ice-cold Hank’s Balanced Salt Solution (HBSS; Thermo Fisher Scientific) supplemented with 10 mM HEPES and penicillin-streptomycin (Thermo Fisher Scientific). Collected tissues were enzymatically digested in 0.05% trypsin and DNase I (Sigma-Aldrich) in the dissociation buffer followed by gentle trituration. Dissociated neurons were seeded in poly-D-lysine-coated culture plates and maintained in serum-free neurobasal medium (Thermo Fisher Scientific) supplemented with B-27 and GlutaMAX (Thermo Fisher Scientific). At 10 days in vitro (DIV), cells were seeded with human P301S tau preformed-fibrils (PFFs; Novus Biologics, NBP2-76794) directly into the medium for 7 days.

**Tau aggregate seeding**

For seeding of tau PFFs (Novus Biologics, NBP2-76794) into the P301S-tau HEK293 cells and the P301L-tau SH-SY5Y cells, Lipofectamine 2000 (Invitrogen, 11668019) was used. 2 μL of Lipofectamine 2000 was used per 1 μg of tau PFFs, and they were separately incubated in Opti-MEM (Gibco, 31984-070) for 5 min. Each solution was then gently mixed and incubated for 15 min before being treated to the cultured cells. For treatment, the seeding mixture was diluted in culture medium at a 1:4 ratio. For seeding of human AD seeds, human brain whole tissue lysate (Novus Biologics, NB820-59363) and human frontal lobe tissue lysate (Novus Biologics) extracted from AD patients were purchased and used. Each lysate was transferred into a tube and incubated with lysis buffer containing 1% sarkosyl and β-mercaptoethanol on an orbital shaker for 1 h at 37°C. The mixture was centrifuged at 150,000 × g for 35 min, and the supernatant was removed. The pellet was resuspended in TBS, aliquoted and stored at -80 °C. Protein concentrations were determined by BCA assay (Thermo Fisher Scientific). The samples were then sonicated at 20% amplitude with 10 pulses. The amount of AD-tau seeds was calculated, and Lipofectamine 2000 was used for seeding into cells.

**Western blotting (WB)**

Cells were harvested using phosphate-buffered saline (PBS; Biosesang, P2007-1) and scrapers and collected by centrifugation at 5000 rpm for 5 min. For non-reducing SDS-PAGE, cell pellets were lysed in Triton X-100 lysis buffer (50 mM Tris, 150 mM NaCl, 100 mM EDTA, 1% Triton X-100, pH 7.6) supplemented with protease and phosphatase inhibitors and incubated on ice for 30 min. The lysates were centrifugated at 5000 rpm for 10 min at 4 °C, and the supernatants were obtained. Protein concentrations were measured using the BCA protein assay kit (Thermo Fisher Scientific, 23225) and diluted to 2 µg/µL. 4× LDS sample buffer (Thermo Fisher Scientific, 84788) was mixed with the protein samples and heated up at 70 °C for 10 min, followed by non-reducing SDS-PAGE. For Triton X-100 fractionation assay, the lysates were centrifugated at 12,000 rpm for 30 min, and the supernatants were taken as the soluble fraction. The remaining pellets were washed with the Triton X-100 buffer by centrifugation at 12,000 rpm for 20 min. The supernatants were discarded, and the insoluble pellets were resuspended in 4× Laemmli sample buffer (Biorad, 161-0747) supplemented with β-mercaptoethanol. The protein samples were subjected to SDS-PAGE. For reducing SDS-PAGE, cells were lysed by RIPA buffer (150 mM NaCl, 1% Triton X-100, 1% sodium deoxycholate, 0.1% Tris–HCl, pH 7.5, 2 mM EDTA) and prepared with 5× protein sample buffer containing β-mercaptoethanol (Elpis Biotech, EBA-1052). After an overnight transfer of the SDS-PAGE gels, the PVDF membranes were blocked with 5% skim milk and washed with PBS-T. The membranes were then incubated with primary antibodies overnight at 4 °C, followed by washing with PBS-T and incubation with horseradish peroxidase (HRP)-conjugated secondary antibodies. Immunoreactive bands were detected using Pierce ECL western blotting substrate (Thermo Fisher Scientific, 32106) and visualized on X-ray films.

**Immunocytochemistry (ICC)**

Cells cultured on cover slips were fixed with 4% paraformaldehyde (PFA) (Biosesang, pc2031-100-00) for 10 min at room temperature, followed by three washes with PBS. The fixed cells were permeabilized with 0.5% Triton X-100 (Sigma Aldrich, T9284) for 10 min and washed with PBS three times. After incubation with 2% BSA blocking solution for 1 h at room temperature, cells were incubated with primary antibodies overnight at 4 °C. After three PBS-washes, cells were incubated with Alexa Fluor-conjugated secondary antibodies for 1 h at room temperature, followed by three 5 min washing with PBS. The slides were then mounted with DAPI-containing mounting medium (Vectashield, H-1500). Images were acquired using LSM700 confocal microscope (Carl Zeiss).

**Immunohistochemistry and immunofluorescence**

Brain tissues were fixed in 4% paraformaldehyde (PFA) at 4 °C for at least 48 h. Following fixation, tissues were washed three times in PBS for 30–40 min each. Coronal sections with 30 μm thickness were prepared using a vibratome (Leica, VT1000S), and the sections were stored at –20 °C in a cryoprotectant buffer containing 2× phosphate buffer, glycerol, and ethylene glycol. For immunohistochemistry, sections were permeabilized with 0.3% Triton X-100 for 20 min at room temperature, treated with 3% hydrogen peroxide for 15 min to quench endogenous peroxidase activity, and blocked in 2% bovine serum albumin (BSA) for 40 min at room temperature. Sections were incubated with primary antibodies overnight at 4 °C, followed by sequential incubation with the linking reagent and labeling reagent from the Ultra-Streptavidin HRP Kit (BioLegend) for 20 min each at room temperature. Colorimetric development was performed with DAB buffer and DAB chromogen, and stained sections were mounted using Vectamount mounting medium (Vector Laboratories, H-5000). Between each step, sections were washed three times in PBS for 5 min. Slides were imaged and analyzed using Axio Scan Z1 slide scanner (Zeiss). For immunofluorescence, the blocked sections were incubated with primary antibodies overnight at 4 °C, followed by secondary antibodies for 40 min at room temperature. Nuclei were counterstained using antifade mounting medium with DAPI (Vector Laboratories). Fluorescent signal detection and image acquisition were performed using Axio Scan Z1 slide scanner (Zeiss) or LSM 700 confocal microscope (Zeiss).

**Proximity ligation assay (PLA)**

PLA was carried out using the Duolink in situ red starter kit (Sigma Aldrich, DUO92101) according to the manufacturer’s protocol. Rat primary hippocampal neurons cultured on glass coverslips were fixed with 4% PFA and permeabilized with 0.5% Triton X-100. After blocking in the provided blocking solution, cells were incubated overnight at 4 °C with anti-p62 (Abcam, ab56416) and anti-Tau5 (Invitrogen, AHB0042) primary antibodies. Subsequent hybridization, ligation, and amplification steps were performed sequentially according to the kit protocol. Nuclei were counterstained with DAPI-containing mounting medium (Vectashield, H-1500), and images of PLA^+^ fluorescence signals were acquired using LSM700 confocal microscope (Carl Zeiss).

**Antibodies**

The following are primary antibodies used: mouse monoclonal anti-SQSTM1/p62 (Abcam, ab56416; 1:20,000 for WB, 1:300 for ICC), rabbit polyclonal anti-LC3 (Sigma Aldrich, L7543; 1:10,000 for WB, 1:300 for ICC), rabbit polyclonal anti-GAPDH (Bioworld, AP0066; 1:5,000 for WB), rabbit polyclonal anti-β-actin (Bioworld, AP0060; 1:20,000 for WB), mouse monoclonal anti-Tau5 (Invitrogen, AHB0042; 1:1000 for WB, 1:100 for ICC), mouse monoclonal anti-T22 (Sigma Aldrich, ABN459; 1:1000 for WB, 1:100 for ICC), mouse monoclonal anti-Tau12 (Sigma Aldrich, MAB2241; 1:1000 for WB), rabbit polyclonal anti-phospho-tau (Ser 199) (Abcam, ab81268; 1:2000 for WB, 1:200 for IHC), rabbit polyclonal anti-phospho-tau (Ser 396) (Invitrogen, 44-752G; 1:2000 for WB), rabbit polyclonal anti-phospho-tau (Ser 214/212) (Thermo Fisher Scientific, MN1060; 1:2000 for WB), mouse monoclonal anti-AT8 (Thermo Fisher Scientific, MN1020; 1:1000 for WB, 1:100 for IHC), mouse monoclonal anti-c-Myc (Santa Cruz Biotechnology, H1721; 1:2,000 for WB), rabbit monoclonal anti-GFAP (Cell Signaling Technology, D1F4Q; 1:500 for IHC), and mouse monoclonal anti-NeuN (Abcam, ab104224; 1:200 for IHC). The following are secondary antibodies used: anti-rabbit IgG-HRP (Cell Signaling Technology, 7074; 1:5,000), anti-mouse IgG-HRP (Cell Signaling Technology, 7076; 1: 5,000), Alexa fluor 488 goat anti-rabbit IgG (A11008; 1:500), and Alexa fluor 555 goat anti-mouse IgG (Invitrogen, A11029; 1:500).

**HTRF ELISA**

Cells lysis was performed following the manufacturer’s instructions. For HTRF analysis, 10 µL of lysate was loaded into a 96-well low-volume plate. Anti-human TAU-d2 and anti-human TAU-Tb^3+^-cryptate conjugates were each diluted 1:50 in diluent and added to the wells. Plates were incubated at 37 °C for 2 h and at room temperature for 10 min. Fluorescence was measured at 665 nm and 620 nm using a Victor Nivo plate reader (Revvity).

**RNA extraction and qRT-PCR analysis**

Total RNA from cells or tissues was extracted using TRizol reagent (Invitrogen, 15596026). cDNA was synthesized from 2 µg of RNA using the PrimeScript 1st strand cDNA synthesis kit (Takara, 6110A). Synthesized cDNA was diluted in distilled water, and 1 µg cDNA was used for quantitative RT-PCR. Gene expression was analyzed using 2 × Fast Q-PCR master mix with SYBR (SMOBIO, TQ1210).

**Animals**

JNPL3 mice (Taconic Biosciences) and rTg4510 mice (KBRI) were maintained in a pathogen-free facility under controlled temperature and humidity conditions with a 12 h light/dark cycle. All animal procedures complied with institutional guidelines and were approved by the Institutional Animal Care and Use Committee (IACUC) of AUTOTAC Bio Inc. and KBRI.

**Brain tissue preparation**

Mice were anesthetized by intraperitoneal injection of 2% avertin (2,2,2-Tribromoethanol) and perfused with 0.9% saline. Extracted brain tissues were stored at –80 °C until use in immunoblotting experiments. To separate soluble and insoluble protein fractions, Triton X-100 lysis buffer (50 mM Tris, 150 mM NaCl, 100 mM EDTA, 1% Triton X-100) supplemented with protease and phosphatase inhibitors was used. Brain tissues were fully homogenized in the Triton X-100 buffer and incubated on ice for 30 min, followed by centrifugation at 13,000 rpm for 20 min at 4 °C. The supernatant (soluble protein fraction) was collected, and protein concentration was determined using the BCA protein assay kit (Thermo Fisher Scientific). The soluble protein sample was prepared in 5× SDS sample buffer containing β-mercaptoethanol, followed by boiling at 100 °C for 5 min. To obtain the Triton X-100 insoluble protein fraction, the remaining pellet was re-suspended in the Triton X-100 buffer and centrifuged again at 13,000 rpm for 15 min at 4 °C. After removing the supernatant, the pellet was resuspended with PBS and prepared in 5× SDS sample buffer, followed by boiling at 100 °C for 5 min. For sarkosyl fractionation, mouse brains were homogenized in extraction buffer (0.8 M NaCl, 10% sucrose, 1 mM EGTA, 50 mM Tris-HCl, and protease and phosphatase inhibitors) and centrifuged at 10,000 × g for 10 min at 4 °C. The pellet was re-extracted and the supernatants were combined. The supernatant was brought to 1% final sarkosyl, incubated for 2 h with gentle mixing, and ultracentrifuged at 100,000 × g for 60 min. Collect the supernatant as sarkosyl-soluble fraction. The pellet was washed once with extraction buffer, ultracentrifuged again at 100,000 × g for 30 min, and resuspended. These fractions were subjected to SDS-PAGE.

**Behavioral tests on mice**

The behavioral tests were performed according to IACUC approval (AUTOTAC Bio Inc., ATB-2104–03-1 and ATB-2209-03-5; KBRI, IACUC-21-00046). For the forelimb grip strength test, a grip strength meter (Bioseb, BIO-GS3) was used. Three consecutive measurements were done for each mouse, and the average was calculated. For the motor coordination test, a rotarod (Bioseb, BX-ROD) was used. Each trial consisted of three attempts. The rotating speed was gradually increased from 4 to 40 rpm in the total time of 300 sec. Open-field testing was used to measure the overall locomotor activity, novelty seeking, and anxiety levels of mice. The mice were placed in a cuboid box without a lid, and their movements were observed and recorded by video. The mice were placed in the center of the open field box (40 × 40 × 40 cm) and moved freely for 30 min.

Novel object recognition test measures hippocampus-dependent memory using the rodent's preference for new objects. In the habituation step (day 1), the mice were placed on the box (40 × 40 × 40 cm) and moved freely for 30 min. In the training step (day 2), two identical objects were put in the box, and the mouse explored them freely for 15 min. In the test stage (day 3), one of two identical objects was replaced with a new object, and the time to explore the new object was measured for 5 min.

Morris water maze tests were performed to evaluate learning and spatial and reference memory. Mice were trained in a circular tank filled with opaque water maintained at 22–24 °C, containing a submerged escape platform hidden 1 cm below the water surface in a fixed location. During training trials, mice were released into the pool from carrying start points and allowed to swim freely for 60 sec. Mice that failed to locate the platform within this time were guided to it and permitted to remain on the platform for 15 sec. Training trials were conducted daily for 6 consecutive days. On day 7, a probe trial was performed in which the platform was removed. Swim paths were recorded, and spatial memory was assessed by quantifying escape latency, time spent in the target quadrant, entries into the target quadrant, and number of crossings over the previous platform location.

Hindlimb clasping behavior was assessed by suspending mice by the base of the tail for several seconds. Posture and hindlimb movement were scored on a 0–5 scale, with 0 indicating normal splaying and 5 indicating severe clasping of both hindlimbs toward the abdomen. For food-motivated maze test, mice were food-restricted to maintain 95% of their baseline body weight for approximately 12 h prior to testing to enhance food-seeking motivation. On the day of testing, mice were first allowed to acclimate to the behavioral testing room for 30 min. Following acclimation, mice were given a 10-min free exploration period inside the maze. After exploration, a single food pellet was placed at the end of one of the arms, and the mice were placed at the starting point. Each mouse was allowed 1 min per trial to locate the baited arm, and the task was repeated for 10 trials. Their behavior was assessed by recording the latency to first correct choice and the number of trials required to reach the second correct choice. The neuromuscular coordination and endurance were assessed using a manual coat-hanging wire apparatus. The hanger was placed 45 cm above the ground. Each mouse was subjected to hanging in the middle of the wire in an upside-down posture, and the time duration was measure throughout three trials. For beam-walking test, a 1-m-long beam was elevated 50 cm above the tabletop with a black goal box at the end. Mice were trained to traverse each beam, and on the test day, the latency to cross 80 cm was recorded over trials.

**EchoMRI body composition**

Body composition was assessed using quantitative magnetic resonance (EchoMRI LLC). Mice were weighed, briefly acclimated to the device room, and placed unanesthetized into the manufacturer’s restrainer tube. Each mouse underwent 2–3 consecutive scans and values for fat mass, lean mass, free water, and total water were exported.

**Quantification and statistical analysis**

Data are presented as mean ± S.E.M. of three independent experiments. Statistical significance was assessed using a two-tailed student’s t-test, with *p< 0.05, **p < 0.01, ***p < 0.001, and ****p < 0.0001 considered statistically significant. MS data were analyzed using Perseus software (version 1.6.15.0). iBAQ intensities were log2-transformed, and proteins with valid values in at least two-thirds of each group were retained. Missing values were imputed from a normal distribution (width = 0.3, down-shift = 1.8). Two-sample t-tests were performed, and a significance criterion of p-value < 0.05 was applied for each group comparison.

**Chemical synthesis and analytical data of AUTOTACS**

^1^H-NMR (Supplementary Fig. X) and ^13^C-NMR (Supplementary Fig. X) spectra were recorded on Bruker Avance III 400 MHz and Bruker Fourier 300 MHz and TMS was used as an internal standard. LC/MS was taken on a quadrupole Mass Spectrometer on Agilent 1260HPLC and 6120MSD (Column: C18 (50 × 4.6 mm, 5 μm) operating in ES (+) or (-) ionization mode; T = 30 °C; flow rate = 1.5 mL/min; detected wavelength: 220 nm, 254nm.

**Scheme 1.** Synthesis of ATB2047

*1.1 Synthesis of ATB2047*

To a solution of **compound** **1-1** (10.0 g, 61.0 mmol, 1.00 eq) and NaH (3.05 g, 76.2 mmol, 1.25 eq, 60% in mineral oil) in dry dimethyl sulfoxide (60 mL) was stirred at 15 °C for 30 min. Then a solution of methyl 3-bromobenzoate (16.4 g, 76.2 mmol, 1.25 eq) in dimethyl sulfoxide (30 mL) was added at 20 °C. The resulting mixture was stirred at 25 °C for 2 hrs. The reaction was quenched with saturated aqueous NH_4_Cl solution. The mixture was poured into water (300 mL) and petroleum ether (100 mL) and stirred for 0.5 h at room temperature. Then filtered to give the **compound 1-2** (17.0 g, crude) as yellow solid. (TLC: PE/EA=10/1, R_f_=0.4)

^1^H-NMR (CDCl_3_, 400 MHz): δ 8.07 (s, 1H), 7.88 (d, *J* = 7.6 Hz, 1H), 7.59-7.69 (m, 2H), 7.46 (s, 1H), 7.30-7.37 (m, 1H), 6.90 (d, *J* = 8 Hz, 1H), 6.69 (s, 1H), 6.07 (s, 2H).

*1. 2 Synthesis of* ***compound 1-3***

To a solution of compound 2 (17.0 g, 49.0 mmol, 1.00 eq) and N_2_H_4_.H_2_O (2.82 g, 56.3 mmol, 1.15 eq) in ethanol (270 mL) was refluxed for 2 hrs. The mixture was cooled to room temperature and filtered to give the compound 3 (13.0 g, 77.4%) as off-white solid. (TLC: PE/EA=5/1, R_f_=0.3)

^1^H-NMR (DMSO_d_6_, 400 MHz): δ 13.08 (s, 1H), 8.02 (s, 1H), 7.82 (d, *J* = 7.6 Hz, 1H), 7.32-7.52 (m, 4H), 7.08 (s, 1H), 6.98 (d, *J* = 8 Hz, 1H), 6.05 (br s, 2H).

*1. 3 Synthesis of* ***compound 1-4***

To a solution of **compound 1-3** (50.0 g, 146 mmol, 1.00 eq) and 2,2'-(ethane-1,2-diylbis(oxy))diethanamine (64.7 g, 437 mmol, 3.00 eq) in 1,4-dioxane (500 mL) was add NaH (17.5 g, 437 mmol, 3.00 eq) slowly at 25 °C. Then added Pd_2_(dba)_3_ (5 g, 5.46 mmol, 0.37 eq) and BINAP (10 g, 16.1 mmol, 9.07 eq) at 25 °C under N_2_. The mixture was stirred overnight at 100 °C under N_2_. The reaction was cooled to room temperature, then quenched with water (1000 mL), extracted with EA (500 mL), and concentrated. The crude was purified by column chromatography (DCM/MeOH=50/1~5/1) to give the **compound 1-4** (26.0 g, 43.4%). (TLC: DCM/MeOH=7/1, R_f_=0.4)

^1^H-NMR (CDCl_3_, 400 MHz): δ 7.29 (s, 2H), 7.20-7.22 (m, 1H), 6.99-7.01(m, 2H), 6.83 (d, *J* = 8 Hz, 1H), 6.69 (s, 1H), 6.58 (d, *J* = 8 Hz, 1H), 5.97 (s, 1H), 3.71-3.73 (m, 2H), 3.55-3.66 (m, 6H), 3.47 (s, 2H), 3.27-3.29 (m, 2H), 2.98-3.01 (m, 2H)

*1. 4 Synthesis of ATB2047*

To a solution of **compound 1-4** (60.0 g, 146 mmol, 1.00 eq) and **compound 1-7** (38.3 g, 146 mmol, 1.00 eq) in MeOH (1000 mL) was stirred overnight at 50 °C. The reaction was concentrated and purified by column chromatography (DCM/MeOH=50/1~20/1) to give the **ATB2047** (10.0 g, 9.99%). (TLC: DCM/MeOH=10/1, R_f_=0.6)

^1^H-NMR (CDCl_3_, 400 MHz): δ7.34-7.38 (m, 2H), 7.19-7.26 (m, 4H), 7.04-7.12 (m, 3H), 6.69 (d, *J* = 7.6 Hz, 1H), 6.81 (d, *J* = 8 Hz, 1H), 6.68 (s, 1H), 6.58-6.61 (m, 1H), 6.53-6.56 (m, 1H), 6.42-6.44 (m, 1H), 6.38-6.41 (m, 1H), 5.96 (s, 2H), 4.90 (s, 2H), 4.53-4.56 (m, 1H), 3.94-3.96 (m, 1H), 3.80-3.91 (m, 1H), 3.78-3.81 (m, 2H), 3.66-3.69 (m, 2H), 3.62 (s, 4H), 3.18-3.38 (m, 6H)

LC/MS [mobile phase: from 90% water (0.05% FA) and 10% CH_3_CN to 5% water (0.1% FA) and 95% CH_3_CN in 6.0 min, finally under these conditions for 0.5 min.] purity is >98% (254 nm), Rt = 2.980 min; Mass Calcd.:684; MS Found: 685 [MS+1].

*1. 5 Synthesis of* ***compound 1-6***

To a solution of **compound 1-6** (50.0 g, 455 mmol, 1.00 eq) and 1-(bromomethyl)-4-fluorobenzene (94.5 g, 500 mmol, 1.10 eq) in DMF (1000 mL) was add K_2_CO_3_ (75.3 g, 545 mmol, 1.2 eq) at 25 °C. The mixture was stirred for 6 h at 25 °C. The reaction was poured into water, then was filtered to give the **compound 1-6** (100 g, crude). (TLC: PE/EA=5/1, R_f_=0.6)

*1. 6 Synthesis of* ***compound 1-7***

To a solution of **compound 1-6** (100 g, 459 mmol, 1.00 eq) in EtOH (2000 mL) was added KOH (33.4 g, 496 mmol, 1.2 eq) and water (200 mL), then added (*R*)-2-(chloromethyl)oxirane (127 g, 1376 mmol, 3 eq) at room temperature. The mixture was stirred overnight at 30 °C. The reaction was quenched by water, extracted with EA, followed by brine and concentrated to give **compound 1-7** (90 g , crude). (TLC: PE/EA=3/1, R_f_=0.6)

**Scheme 2.** Synthesis of **ATB2055**

*2.1 Synthesis of* ***compound 2-2***

To a solution of **compound** **2-1** (10.0 g, 61.0 mmol, 1.00 eq) in dry dimethyl sulfoxide (100 mL) was added NaH (3.05 g, 76.2 mmol, 1.25 eq, 60% in mineral oil) in portions, the mixture was stirred at room temperature for 30 min. Then a solution of methyl 3-bromo-5-iodobenzoate (26.0 g, 76.2 mmol, 1.25 eq) in dimethyl sulfoxide (30 mL) was added dropwise, the resulting mixture was stirred at room temperature for 2 hrs. The reaction was quenched with saturated aqueous NH_4_Cl solution. The mixture was poured into water (300 mL) and petroleum ether (100 mL) and stirred for 0.5 h at room temperature. Then filtered to give **compound 2-2** (15.0 g, crude) as yellow solid. (TLC: PE/EA=3/1, R_f_=0.7)

*2.2 Synthesis of* ***compound 2-3***

To a solution of **compound 2-2** (15.0 g, 31.7 mmol, 1.00 eq) in ethanol (200 mL) was added N_2_H_4_.H_2_O (1.82 g, 36.5 mmol, 1.15 eq), the mixture was stirred at reflux for 2 h. The mixture was cooled to room temperature and filtered to give **compound 2-3** (12.0 g, 80.5%) as yellow solid. (TLC: PE/EA=2/1, R_f_=0.5)

*2.3 Synthesis of* ***compound 2-4***

To a solution of **compound 2-3** (4.0 g, 8.53 mmol, 1.00 eq) and 2,2'-(ethane-1,2-diylbis(oxy))diethanamine (3.79 g, 25.6 mmol, 3.00 eq) in 1,4-dioxane (50 mL) was add NaH (1.02 g, 25.6 mmol, 3.00 eq) slowly at room temperature. Then added Pd_2_(dba)_3_ (0.4 g, 0.44 mmol, 0.05 eq) and BINAP (0.82 g, 1.32 mmol, 0.15 eq) under N_2_. The mixture was stirred overnight at 100 °C under N_2_. The reaction was cooled to room temperature, then quenched with water (100 mL), extracted with EtOAc (100 mL × 2), and concentrated. The crude was purified by column chromatography (DCM/MeOH=50/1~10/1) to give **compound 2-4** (1.0 g, 24.0%) as yellow solid (TLC: DCM/MeOH=10/1, R_f_=0.2)

*2.4 Synthesis of* ***ATB2055***

To a solution of **compound 2-4** (3.0 g, 6.13 mmol, 1.0 eq) in 1,2-dichloroethane (30 mL) was added 3-(4-fluorobenzyloxy)benzaldehyde (1.2 g, 5.22 mmol, 0.85 eq), the mixture was stirred at room temperature overnight. NaBH(OAc)_3_ (1.3 g, 6.13 mmol, 1.0 eq) was added in portions. The mixture was stirred at room temperature for 30 min. Poured into 50 mL water, extracted with DCM (50mL × 2), the DCM layer was dried over Na_2_SO_4_ and concentrated, the residue was purified by column chromatography (DCM/MeOH=50/1~10/1) to give **ATB2055** (1.1 g, 25.5%) as white solid. (TLC: DCM/MeOH=10/1, R_f_=0.3)

^1^H-NMR (CDCl_3_, 400 MHz): δ 7.291-7.327 (m, 2H), 7.162-7.200 (m, 3H), 7.100 (t, J=1.2Hz, 1H), 6.970-7.038 (m, 3H), 6.892 (d, J=7.6Hz, 1H), 6.789-6.826 (m, 3H), 6.649 (t, J=1.6Hz, 1H), 6.627(s, 1H), 5.962 (s, 2H), 4.891 (s, 2H), 4.414(t, J=5.6Hz, 1H), 3.813 (s, 2H), 3.573-3.626 (m, 8H), 3.153-3.190 (m, 2H), 2.862 (t, J=4.2Hz, 2H).

LC/MS [mobile phase: from 90% (0.05%FA in water) and 10% (0.05%FA in ACN:H2O=9:1 to 5% (0.05%FA in water) and 95% (0.05%FA in ACN:H2O=9:1) in 6.0 min, finally under these conditions for 0.5 min.] purity is 97.4 %, Rt = 3.165 min; MS Calcd.:702.2; MS Found: 352.2([M/2+1]^+^), 703.2([M+1]^+^).

**Scheme 3.** Synthesis of **ATB2056**

*3.1 Synthesis of* ***intermediate***

To a solution of concentrated sulfuric acid (12 mL) in water (76 mL) was added methyl 3-amino-5-bromobenzoate (20.0 g, 87.0 mmol, 1.0 eq) at 0 °C, then a solution sodium nitrite (6.0 g, 87.0 mmol, 1.0 eq) was added drop wise while the solution turned to yellow. After additional 30 min the solution was heated and refluxed for 30 min. After cooling to room temperature. the brownish solution was extracted four times with DCM. The combined organic phase was dried over sodium sulfate and concentrated under reduced pressure. Then the residue was purified by column chromatography on silica gel to give intermediate (5.0 g, 24.9%) as a red solid. (TLC: EA/PE=1:2, Rf=0.7)

*3.2 Synthesis of* ***compound 3-2***

To a solution of **compound** **3-1** (25.0 g, 200 mmol, 1.00 eq) and 1-(bromomethyl)-4- fluorobenzene (38.7 g, 200 mmol, 1.00 eq) in DMF (200 mL) was added K_2_CO_3_ (69.0 g, 500 mmol, 2.50 eq), the mixture was stirred at 100 °C overnight. Cooled to room temperature, the mixture was poured into ice water, the solid was filtered and washed with water, dried in vacuum to give **compound 3-2** (25.0 g, 53.0%) as yellow solid. (TLC: PE/EA=3/1, R_f_=0.7)

*3.3 Synthesis of* ***compound 3-3***

To a mixture of hydroxylamine hydrochloride (9.2 g, 130.4 mmol, 2.0 eq) and NaOH (82 mL, 1.6 M in H_2_O) in MeOH (82 mL) was added **compound 3-2** (15.0 g, 65.2 mmol, 1.0 eq). The mixture was stirred at 70 °C for 1 h. MeOH was evaporated in vacuum and the solution was extracted with EA. The organic layer was dried with anhydrous sodium sulfate and evaporated in vacuum to provide **compound 3-3** (crude) as a yellow solid. (TLC: PE/EA=3/1, Rf=0.7)

*3.4 Synthesis of* ***compound 3-4***

To a stirred solution of **compound 3-3** (crude) in acetic acid (160 mL) was added zinc powder (21.2 g, 326.1 mmol, 5.0 eq) in 6 portions under 70 °C. After being stirred for 1 h at 70 °C, the reaction mixture was filtered and the most solvent was evaporated in a vacuum. Excess of ammonia was added to the solution and the solution was extracted with EA. The organic layer was washed with water and dried with Na_2_SO_4_. After filtration and evaportation under reduced pressure, the residue was purified by column chromatography on silica gel to give **compound 3-4** (10.0 g, 66.4 %) as white solid. (TLC: DCM/MeOH=10:1, Rf=0.4).

*3.5 Synthesis of* ***compound 3-5***

A mixture of **compound 3-4** (10.0 g, 43.3 mmol, 1.3 eq), 2-(2-(2-hydroxyethoxy)ethoxy)ethyl 4-methylbenzenesulfonat**e** (10.0 g, 32.9 mmol, 1.0 eq), K_2_CO_3_ (9.0 g, 65.8 mmol, 2 eq) and NaI (2.0 g, 13.2 mmol, 0.4 eq) in ACN (500 mL) was stirred at 90 °C for 15 h and cooled to room temperature. To the mixture was added (Boc)_2_O (18.0 g, 82.3 mmol, 2.5 eq) and stirred at room temperature for 5 h. The mixture was filtered and concentrated in vacuum, the residue was purified by column chromatography on silica gel to give **compound 3-5** (6.0 g, 39.5 %) as yellow oil. (TLC: PE/EA=1:1, Rf=0.3)

*3.6 Synthesis of* ***compound 3-6***

A solution of **compound 3-5** (5.0 g, 10.8 mmol, 1.0 eq) in DCM (125 mL) was added Ag_2_O (3.7 g, 16.2 mmol, 1.5 eq), NaI (1.8 g, 11.9 mmol, 1.1 eq) and TsCl (2.7 g, 14.0 mmol, 1.3 eq) at 0 °C. The reaction mixture was then allowed to warm to room temperature and stirred for 1 h. The mixture was filtered through celite and the filtrate was washed with 10% aq. NaHCO_3_. The organic layer was dried over anhydrous sodium sulfate and filtered. The filtrate was evaporated under reduced pressure and the crude product was purified by column chromatography on silica gel to give **compound 3-6** (6.0 g, 90.0 %) as colorless oil. (TLC: EA/PE=1/2, Rf=0.4)

*3.7 Synthesis of* ***compound 3-7***

A mixture of methyl 3-bromo-5-hydroxybenzoate (2.2 g, 9.7 mmol, 1.0 eq), **compound 3-6** (6.0 g, 9.7 mmol, 1.0eq), K_2_CO_3_ (6.7 g, 48.6 mmol, 5.0 eq) and NaI (0.6 g, 3.9 mmol, 0.4 eq) in ACN (360 mL) was stirred at 90 °C for 16 h. Cooled to room temperature, the mixture was filtered and concentrated in vacuum, the residue was purified by column chromatography on silica gel to give **compound 3-7** (5.0 g, 76.2 %) as red oil. (TLC: EA/PE=1/2, Rf=0.5)

*3.8 Synthesis of* ***compound 3-8***

To a solution of 1-(benzo[d][1,3]dioxol-5-yl)ethanone (1.1 g, 6.7 mmol, 1.0 eq) in dry dimethyl sulfoxide (22 mL) was added NaH (0.3 g, 8.1 mmol, 1.2 eq, 60% in mineral oil) in portions, the mixture was stirred at room temperature for 30 min. Then a solution of **compound 3-7** (5.5 g, 8.1 mmol, 1.2 eq) in dimethyl sulfoxide (11 mL) was added dropwise, the resulting mixture was stirred at room temperature for 2 h. The reaction was quenched with saturated aqueous NH_4_Cl solution, extracted with EA, The organic layer was washed with water and dried with Na_2_SO_4_. After filtration and evaportation under reduced pressure, the residue was purified by column chromatography on silica gel to give **compound 3-8** (4.2 g, 63.9 %) as a red solid. (TLC: EA/PE=1:2, Rf=0.4).

*3.9 Synthesis of* ***compound 3-9***

To a solution of **compound 3-8** (4.2 g, 5.2 mmol, 1.00 eq) in ethanol (65 mL) was added N_2_H_4_.H_2_O (0.3 g, 6.0 mmol, 1.15 eq), the mixture was stirred at reflux for 2 hrs. The mixture was cooled to room temperature and evaporated under reduced pressure, the residue was purified by column chromatography on silica gel to give **compound 3-9** (2.9 g, 69.4 %) as a red solid. (TLC: EA/PE=1:1, Rf=0.3).

*3.10 Synthesis of* ***ATB2056***

To a solution of TFA (3 mL) in DCM (30 mL) was added **compound 3-9** (2.9 g, 3.6 mmol, 1.0 eq), the mixture was stirred at r.t. for 30 min. Concentrated under reduced pressure, the residue was dissolved in EA, washed with aqueous NaHCO_3_ and brine, dried over Na_2_SO_4_, filtered and concentrated, the residue was purified by prep-HPLC and freeze dried to give **ATB2056** (1.2 g, 47.2%) as a white solid. (TLC: DCM/MeOH=10:1, Rf=0.3)

^1^H-NMR (DMSO_d_6_, 400 MHz): δ 13.306 (br, 1H), 7.603 (t, J=1.2Hz, 1H), 7.463-7.499 (m, 2H), 7.401-7.409 (m, 1H), 7.371 (d, J=1.6Hz, 1H), 7.327 (d, J=8.0Hz, 1H), 7.179-7.238 (m, 4H), 7.111 (s, 1H), 7.012 (d, 1H, J=8.0Hz), 6.980 (s, 1H), 6.885 (d, J=7.6Hz, 1H), 6.845 (dd, J=8.0Hz, J=2.4Hz, 1H), 6.070 (s, 2H), 5.045 (s, 2H), 4.174 (t, J=4.0Hz, 2H), 3.761 (t, J=4.8Hz, 2H), 3.681 (s, 2H), 3.597-3.620 (m, 2H), 3.524-3.546 (m, 2H), 3.483 (t, J=6.4Hz, 2H), 2.627 (t, J=5.6Hz, 2H).

^13^C-NMR (DMSO_d_6_, 400 MHz): δ 163.381, 160.959, 160.232, 158.711, 148.255, 147.506, 142.997, 133.882, 133.851, 130.358, 130.282, 129.579, 123.070, 120.861, 120.465, 119.319, 116.783, 115.761, 115.549, 114.709, 113.299, 111.026, 109.158, 106.007, 101.671, 100.529, 70.524, 70.384, 70.132, 69.297, 68.793, 68.156, 53.193, 48.395.

LC/MS [mobile phase: from 90% (0.05%FA in water) and 10% (0.05%FA in ACN:H2O=9:1 to 5% (0.05%FA in water) and 95% (0.05%FA in ACN:H2O=9:1) in 6.0 min, finally under these conditions for 0.5 min.] purity is 99.8 %, Rt = 3.612 min; MS Calcd.:703.2; MS Found: 352.6([M/2+1]^+^), 704.1([M+1]^+^).

**Scheme 4.** Synthesis of **ATB2082**

*4.1 Synthesis of* ***core 4-1-2***

To a solution of **core 4-1-1** (10.0 g, 81.9 mmol, 1.0 eq), 1-(bromomethyl)-4-fluorobenzene (17.0 g, 90.0 mmol, 1.1 eq), K_2_CO_3_ (13.6 g, 98.2 mmol, 1.2 eq) and KI (1.36 g, 8.19 mmol, 0.1 eq) in DMF (150 mL). The mixture was stirred at 60 °C for 16 hours. The mixture was poured into water (500 mL) filtered and dried to give the core 1-2 (12 g, 63.49%) as an off-white solid. (TLC: DCM/MeOH=10/1, R_f_=0.3)

^1^H-NMR (CDCl_3_, 400 MHz): δ 9.98 (s, 1H), 7.49 - 7.40 (m, 5H), 7.27 - 7.24 (m, 1H), 7.09 (t, *J* = 8.6 Hz, 1H), 5.09 (s, 1H).

*4.2 Synthesis of* ***core 4-1-3***

To a solution of NH_2_OH.HCl (3.63 g, 52.2 mmol. 2.0 eq) in EtOH (35 mL) and 1.6 M NaOH (19 mL) was added **core 4-1**-**2** (6 g, 26.1 mmol, 1.0 eq). The reaction mixture was stirred at 70℃ for 1 hr. The mixture was concentrated in vacuo and added water (20 mL), filtered and dried to give the **core** **4-1**-**3** (5.0 g, 76.63%) as an off-white solid. (TLC: DCM/MeOH=10/1, R_f_=0.3)

*4.3 Synthesis of* ***core 4-1***

To a solution of **core** **4-1**-**3** (5 g, 20.4 mmol, 1.0 eq) in CH_3_COOH (50 mL) was added Zn powder (6.67 g, 0.102 mol, 5.0 eq) at 70 °C. The resulting mixture was stirred at 70℃ for 1 hour under N_2_. The mixture was filtered and the filtration was evaporated to obtain the **core** **4-1** (2.5 g, 49.34%) as an off-white solid. (TLC: DCM/MeOH=15/1, R_f_=0.4)

*4.4 Synthesis of* ***core 4-2-2***

To a solution of **core** **4-2**-**1** (10 g, 0.0609 mol, 1.0 eq) and NaH (1.83 g, 0.0761 mol, 1.25 eq) in DMSO (60 mL) was stirred at 15 °C for 30 min. Then a solution of methyl 3-bromobenzoate (16.4 g, 0.0761 mol, 1.25 eq) in DMSO (30 mL) was added at 25 °C. The resulting mixture was stirred at 25 °C for 2 hrs. The mixture was poured into water (200 mL) and extracted with EA (50 mL × 3). The EA layer was dried over Na_2_SO_4_ filtered and concentrated in vacuo to give the crude **core** **4-2**-**2** (20 g, 69.46%) as a yellow solid. (TLC: DCM/MeOH=20/1, R_f_=0.3)

*4.5 Synthesis of* ***core 4-2***

To a solution of **core** **4-2**-**2** (20 g, 0.0576 mol, 1.0 eq) in EtOH (300 mL) was added hydrazine monohydrate (3.35 g, 0.0662 mol, 1.15 eq). The mixture was at 80℃ for 2 hours. The mixture was filtered and dried to give the **core** **4-2** (8.0 g, 39.24 %) as an off-white solid. (TLC: DCM/MeOH=10/1, R_f_=0.3)

^1^H-NMR (DMSO_d_6_, 400 MHz): δ 13.33 (s, 1H), 8.02 (s, 1H), 7.88 - 7.77 (m, 1H), 7.59 - 7.48 (m, 1H), 7.47 - 7.29 (m, 3H), 7.22 (d, *J* = 11.8 Hz, 1H), 7.06 - 6.97 (m, 1H), 6.07 (d, *J* = 12.1 Hz, 2H).

*4.6 Synthesis of* ***compound 4-2a***

To a solution of oxalyl chloride (1.14 g, 9.0 mmol, 2.0 eq) in DCM (5 mL) cooled to -78 °C was added DMSO (1.40 g, 18.0 mmol, 4.0 eq) in DCM (5 mL) and stirred under nitrogen at -78℃ for 0.5 hour. Then **compound** **4-1a** (1 g, 4.5 mmol, 1.0 eq) in DCM (5 mL) was added and stirred under N_2_ at -78℃ for 0.5 hour. Finally, TEA (3.64 g, 36.0 mmol, 8.0 eq) was added and the reaction mixture was stirred under N_2_ at 25℃ for 1 hour. Another batch was carried out as the above procedure. The mixture was added sat. NaHCO_3_ solution (10 mL), water (20 mL) and extracted with DCM (15 mL × 3). The organic layer was separated and washed with water (10 mL) and brine (10 mL), dried over anhydrous Na_2_SO_4_ and concentrated in vacuo to give the crude **compound** **4-2a** (2.0 g, 80.22 %) as orange oil. (TLC: DCM/MeOH=15/1, R_f_=0.4)

*4.7 Synthesis of* ***compound 4-3a***

To a solution of **core** **4-2** (5 g, 14.6 mmol, 1.0 eq) in DMSO (50 mL) was added diphenylmethanimine (3.97 g, 21.9 mmol, 1.5 eq), Pd_2_(dba)_3_ (2.67 g, 2.92 mmol, 0.2 eq), xantphos (3.38 g, 5.84 mmol, 0.4 eq) and Cs_2_CO_3_ (9.51 g, 29.2 mmol, 2.0 eq). The reaction mixture was stirred under N_2_ at 100 °C for 16 hours. The mixture was cooled to room temperature, added water (60 mL) and extracted with EA (30 mL × 3). The organic layer was washed with water (20 mL) and brine (20 mL), dried over anhydrous Na_2_SO_4_ and concentrated in vacuo. The crude was purified by column chromatography (PE/EtOAc =10/1~1/1) to give the **compound** **4-3a** (2.3 g, 21.02%) as a brown solid. (TLC: PE/EtOAc =2/1, R_f_=0.4)

^1^H-NMR (DMSO_d_6_, 400 MHz): δ 13.14 (s, 1H), 7.69 (d, *J* = 7.2 Hz, 2H), 7.57 - 7.46 (m, 3H), 7.39 - 7.26 (m, 6H), 7.21 - 7.18 (m, 6H), 7.05 - 6.95 (m, 2H), 6.61 (d, *J* = 7.5 Hz, 1H), 6.05 (d, *J* = 10.3 Hz, 2H).

*4.8 Synthesis of* ***compound 4-4a***

To a solution of **compound** **4-3a** (2.3 g, 5.2 mmol, 1.0 eq) in THF (20 mL) was added 3N HCl (15 mL). The mixture was at 25 °C for 3 hours. The mixture was adjusted pH=8 with sat. NaHCO_3_ solution and extracted with EA (20 mL × 3) and the organic layer was washed with water (10 mL) and brine (10 mL), dried over anhydrous Na_2_SO_4_ and concentrated in vacuo. The crude was purified by column chromatography (PE/EtOAc =5/1~1/1) to give the **compound** **4-4a** (800 mg, 55.77%) as a brown solid. (TLC: PE/EtOAc =1/1, R_f_=0.3)

^1^HNMR (DMSO_d_6_, 400 MHz): δ 13.08 (s, 1H), 7.38 - 7.34 (m, 2H), 7.09 - 6.90 (m, 5H), 6.55 (s, 1H), 6.05 (s, 2H), 5.13 (d, *J* = 37.7 Hz, 2H).

*4.9 Synthesis of* ***compound 4-5a***

To a solution of **compound** **4-4a** (350 mg, 1.25 mmol, 1.0 eq) in DCE (10 mL) was added **compound** **4-2a** (410 mg, 1.88 mmol, 1.5 eq), (CH_3_COO)_3_BHNa (531 mg, 2.51 mmol, 2.0 eq) and AcOH (226 mg, 3.76 mmol, 3.0 eq). The reaction mixture was stirred under N_2_ at 25 °C for 16 hours. Another batch was carried out as the above procedure. The mixture was adjusted pH=8 with sat. NaHCO_3_ solution and extracted with DCM (10 mL × 3) and the organic layer was washed with water (5 mL) and brine (5 mL), dried over anhydrous Na_2_SO_4_ and concentrated in vacuo. The crude was purified by column chromatography (PE/EtOAc =5/1~1/1) to give the **compound** **5a** (460 mg, 33.76%) as a brown solid. (TLC: PE/EtOAc =3/1, R_f_=0.4)

^1^H-NMR (DMSO_d_6_, 400 MHz): δ 13.09 (d, *J* = 14.9 Hz, 1H), 7.43 - 7.27 (m, 2H), 7.17 - 6.93 (m, 5H), 6.64 - 6.52 (m, 1H), 6.06 (d, *J* = 9.7 Hz, 2H), 5.61 (d, *J* = 37.1 Hz, 1H), 4.01 - 3.97 (m, 2H), 3.64 - 3.53 (m, 6H), 3.26 (s, 2H), 1.41 (s, 9H).

*4.10 Synthesis of* ***compound 4-6a***

To a solution of **compound** **4-5a** (440 mg, 0.9134 mmol, 1.0 eq) in DCM (4 mL) was added TFA (2 mL). The reaction mixture was stirred at 25℃ for 4 hours. The reaction mixture was evaporated in vacuo to give the crude **compound** **4-6a** (420 mg, 79.71 %) as a brown solid. (TLC: DCM/MeOH=20/1, R_f_=0.3)

*4.11 Synthesis of* ***ATB2082***

To a solution of **compound** **4-6a** (420 mg, 0.987 mmol, 1.0 eq) in DCM (5 mL) was added **core** **4-1** (456 mg, 1.97 mmol, 2.0 eq), EDCI (284 mg, 1.48 mmol, 1.5 eq), HOBT (200 mg, 1.48 mmol, 1.5 eq) and TEA (299 mg, 2.96 mmol, 3.0 eq). The reaction mixture was stirred under N_2_ at 25℃ for 16 hours. The mixture was added water (15 mL) and extracted with DCM (3 mL × 3). The organic layer was separated and washed with water (3 mL) and brine (3 mL), dried over anhydrous Na_2_SO_4_ and concentrated in vacuo. The crude was purified by column chromatography (DCM/MeOH=100/1~50/1) to give **ATB2082** (121 mg, 18.88%) as an off-white solid. (TLC: DCM/MeOH=20/1, R_f_=0.4)

^1^H-NMR (DMSO_d_6_, 400 MHz): δ 13.10 (s, 1H), 8.24 (t, *J* = 6.1 Hz, 1H), 7.49 - 7.45 (m, 2H), 7.38 - 7.34 (m, 2H), 7.23 - 7.17 (m, 3H), 7.14 - 6.91 (m, 5H), 6.90 - 6.81 (m, 3H), 6.57 (s, 1H), 6.05 (s, 2H), 5.61 (d, *J* = 29.8 Hz, 1H), 5.03 (s, 2H), 4.28 (d, *J* = 6.2 Hz, 2H), 3.97 (s, 2H), 3.66 - 3.57 (m, 6H), 3.22 (s, 2H).

LC/MS [mobile phase: from 90% water (0.05%FA) and 10% CH_3_CN to 5% water (0.05% FA) and 95% CH_3_CN in 6.0 min, finally under these conditions for 0.5 min.] purity is >99.0% (254 nm), Rt = 3.712 min; Mass Calcd.:638; MS Found: 639 [MS+1]

**Scheme 5.** Synthesis of **ATB2088**

*5.1 Synthesis of* ***compound 5-A***

To a solution of **core 5-2** (5.00 g, 14.6 mmol, 1.0 eq), ethane-1,2-diamine (17.6 g, 292 mmol, 20 eq), potassium tert-butoxide (4.91 g, 43.8 mmol, 3.0 eq) and BINAP (2.73 g, 4.38 mmol, 0.3 eq) in 1,4-dioxane (50 mL) was added Pd_2_(dba)_3_ (2.67 g, 2.92 mmol, 0.2 eq) at 25 °C under N_2_. The mixture was stirred for 16 hours at 100 °C. The reaction was cooled to room temperature, then filtered. The filtrate was poured into water (150 mL), extracted with DCM (50 mL x 3), washed with water (100 mL), dried over Na_2_SO_4_, filtered and concentrated in vacuo to give the crude product. The crude was purified by column chromatography (DCM/MeOH=50/1~5/1) to give the **compound 5-A** (2.00 g, 42.6%). (TLC: DCM/MeOH=10/1, R_f_=0.4)

^1^H-NMR (DMSO_d_6_, 400 MHz): *δ* 7.41 (d, *J* = 1.5 Hz, 1H), 7.36 (dd, *J* = 8.1, 1.5 Hz, 1H), 7.15 (t, *J* = 7.8 Hz, 1H), 7.05 (s, 1H), 7.00 (d, *J* = 9.6 Hz, 3H), 6.59 (dd, *J* = 8.1, 1.4 Hz, 1H), 6.07 (s, 2H), 5.80 (t, *J* = 5.4 Hz, 1H), 3.17 (dd, *J* = 11.9, 6.0 Hz, 2H), 2.83 (t, *J* = 6.3 Hz, 2H).

*5.2 Synthesis of* ***compound 5-2***

To a solution of **compound 5-1** (5.00 g, 21.7 mmol, 1.0 eq) in EtOH (50 mL) was added 2-(2-aminoethoxy)ethanol (4.56 g, 43.4 mmol, 2.0 eq) and NaBH_4_ (1.23 g, 32.6 mmol, 1.5 eq) at 25 ^o^C. The mixture was stirred for 16 hours at 25 °C. The reaction was quenched with saturated aqueous NH_4_Cl solution (200 mL), extracted with ethyl acetate (50 mL × 3), washed with water (100 mL), dried over Na_2_SO_4_, filtered and concentrated in vacuo to give the crude product. The crude was purified by column chromatography (DCM/MeOH=50/1~5/1) to give the **compound 5-2** (1.50 g, 21.7%). (TLC: DCM/MeOH=10/1, R_f_=3)

^1^H-NMR (CDCl_3_, 400 MHz): *δ* 7.41 (dd, *J* = 8.2, 5.6 Hz, 2H), 7.27 (s, 0.4H), 7.25 - 7.23 (m, 0.7H), 7.07 (t, *J* = 8.6 Hz, 2H), 7.00 (s, 1H), 6.94 (d, *J* = 7.5 Hz, 1H), 6.90 - 6.83 (m, 1H), 5.03 (s, 2H), 3.83 (s, 2H), 3.78 - 3.69 (m, 2H), 3.69 - 3.62 (m, 2H), 3.61 - 3.54 (m, 2H), 2.89 - 2.76 (m, 2H).

*5.3 Synthesis of* ***compound 5-3***

To a solution of **compound 5-2** (1.50 g, 4.70 mmol, 1.0 eq) in DCM (15 mL) was added (Boc)_2_O (1.54 g, 7.05 mmol, 1.5 eq) and TEA (1.43 mg, 14.1 mmol, 3.0 eq) at 0 °C. The mixture was stirred for 4 hours at 25 °C. The mixture was quenched with water (50 mL), extracted with DCM (10 mL × 2), washed with water (10 mL), dried over Na_2_SO_4_, filtered and concentrated in vacuo to give the crude product. The crude was purified by column chromatography (DCM/MeOH=50/1~5/1) to give the **compound 5-3** (1.8 g, 91.4%). (TLC: DCM/MeOH=10/1, R_f_=0.4)

*5.4 Synthesis of* ***compound 5-4***

To a solution of **compound 5-3** (2.00 g, 4.80 mmol, 1.0 eq) in DCM (20 mL) was added NMO (1.12 g, 9.60 mmol, 2.0 eq) and TPAP (170 mg, 0.480 mmol, 0.1 eq). The mixture was stirred for 2 hours at 25 °C. The reaction was quenched with water (50 mL), extracted with DCM (20 mL × 2), washed with water (10 mL), dried over Na_2_SO_4_, filtered and concentrated in vacuo to give the crude product. The crude was purified by column chromatography (DCM/MeOH=50/1~5/1) to give the **compound 5-4** (0.8 g, 40.0%). (TLC: DCM/MeOH=10/1, R_f_=0.6)

^1^H-NMR (CDCl_3_, 400 MHz): *δ* 9.66 (s, 1H), 7.45 - 7.36 (m, 2H), 7.26 - 7.17 (m, 1H), 7.07 (t, *J* = 8.6 Hz, 2H), 6.85 (d, *J* = 8.6 Hz, 3H), 5.01 (s, 2H), 4.50 (s, 1H), 4.10 – 3.95 (m, 2H), 3.80 - 3.25 (m, 6H), 1.53 – 1.30(m, 9H).

*5.5 Synthesis of* ***ATB2088***

To a solution of **compound 5-A** (1.23 g, 3.80 mmol, 1.0 eq) and **compound 5-4** (0.80 g, 1.90 mmol, 0.5 eq) in MeOH (15 mL) was added NaBH(OAc)_3_ (4.03 g, 0.019 mmol, 5.0 eq). The mixture was stirred for 16 hours at 25 ^o^C. The reaction was quenched with NaHCO_3_ saturated solution (5 mL × 3), washed with water (10 mL), dried over Na_2_SO_4_, filtered and concentrated in vacuo to give the crude product. The crude was dissolved in dioxane (10 mL) and the solution was cooled to 0 °C. 4 N HCl in dioxane (10 mL) was added and then stirred for 1 hour at 25 °C. The mixture was concentrated to give crude product. The residue was purified by pre-HPLC (Columns: sunfire 5μm 19-150 mm; Mobile Phase: ACN-H_2_O (0.1%FA); Gradient: 10-60-8GT-400VL) to give the TFA salt solution of product. Then the solution was adjusted the pH value to 8 with NaHCO_3_ saturated solution, extracted with DCM (5 mL × 4) washed with water (5 mL), dried over Na_2_SO_4_, filtered and concentrated in vacuo to product. The residue was lyophilized to give **ATB2088** (100 mg, 4.22%) as yellow oil. (TLC: DCM/MeOH=10/1, R_f_=1)

^1^H-NMR (DMSO_d_6_, 400 MHz): *δ* 13.10 (s, 1H), 7.48 (dd, *J* = 8.5, 5.7 Hz, 2H), 7.38 (d, *J* = 1.4 Hz, 1H), 7.33 (d, *J* = 7.8 Hz, 1H), 7.20 (dd, *J* = 12.3, 5.6 Hz, 3H), 7.11 (t, *J* = 7.8 Hz, 1H), 6.97 (s, 5H), 6.89 (d, *J* = 7.5 Hz, 1H), 6.84 (dd, *J* = 8.2, 2.0 Hz, 1H), 6.55 (d, *J* = 7.3 Hz, 1H), 6.05 (s, 2H), 5.61 (s, 1H), 5.04 (s, 2H), 3.67 (s, 2H), 3.46 (dd, *J* = 8.9, 5.5 Hz, 4H), 3.15 (s, 3H), 2.77 (t, *J* = 6.1 Hz, 2H), 2.72 (t, *J* = 5.5 Hz, 2H), 2.61 (t, *J* = 5.6 Hz, 2H).

LC/MS [mobile phase: from 90% A (0.05%FA in water) and 10% B (0.05%FA in CAN/H_2_O=9/1）to 5% A (0.05%FA in water) and 95% B (0.05%FA in CAN/H_2_O=9/1) in 0 min, finally under these conditions for 0.5 min.] purity is >99.0% (220 nm), Rt = 2.118 min; Mass Calcd.:623; MS Found: 624 [MS+1].

**^1^H-NMR spectrum of AUTOTACS used in this study**

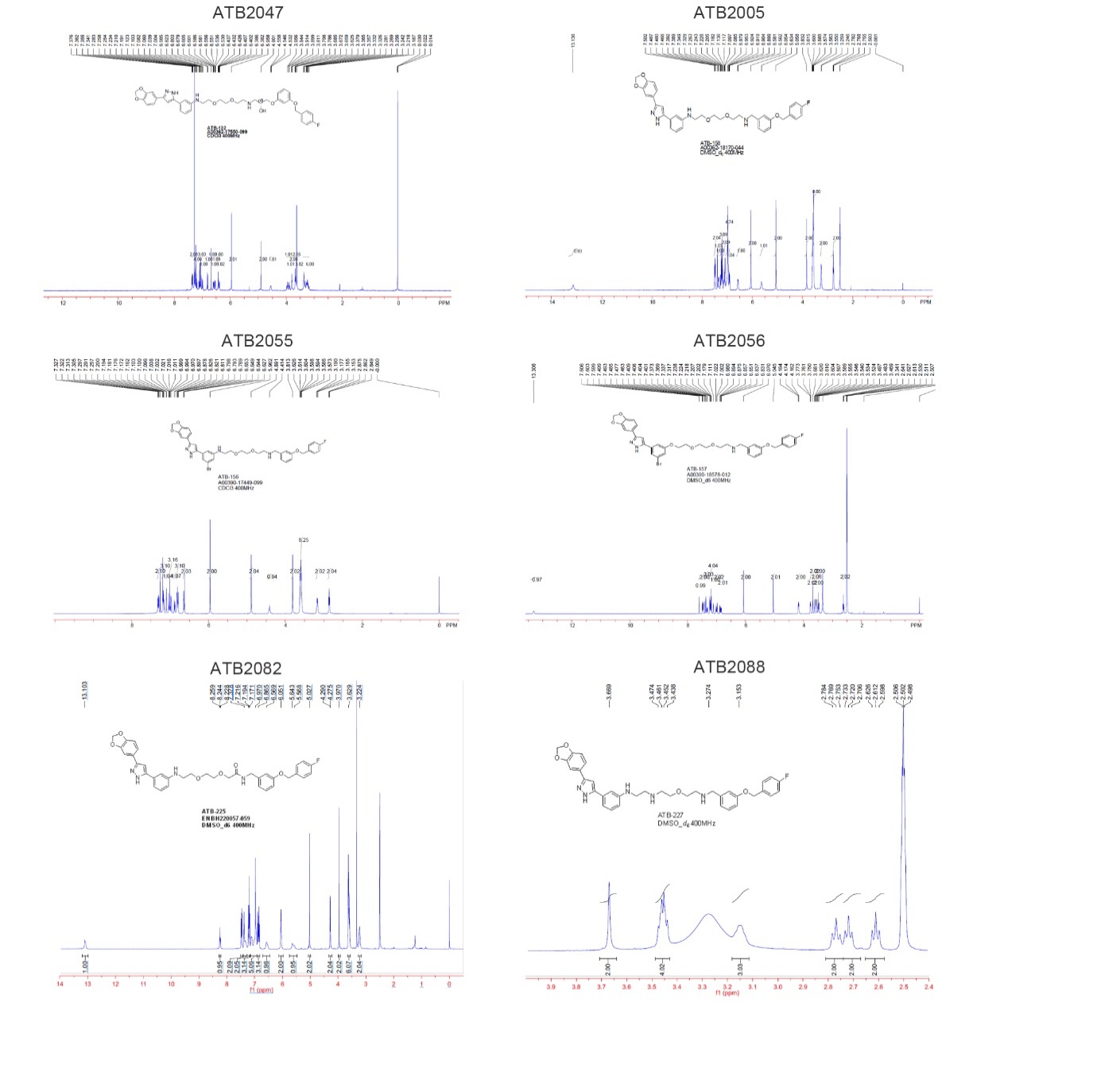

**HPLC data of AUTOTACS used in this study**

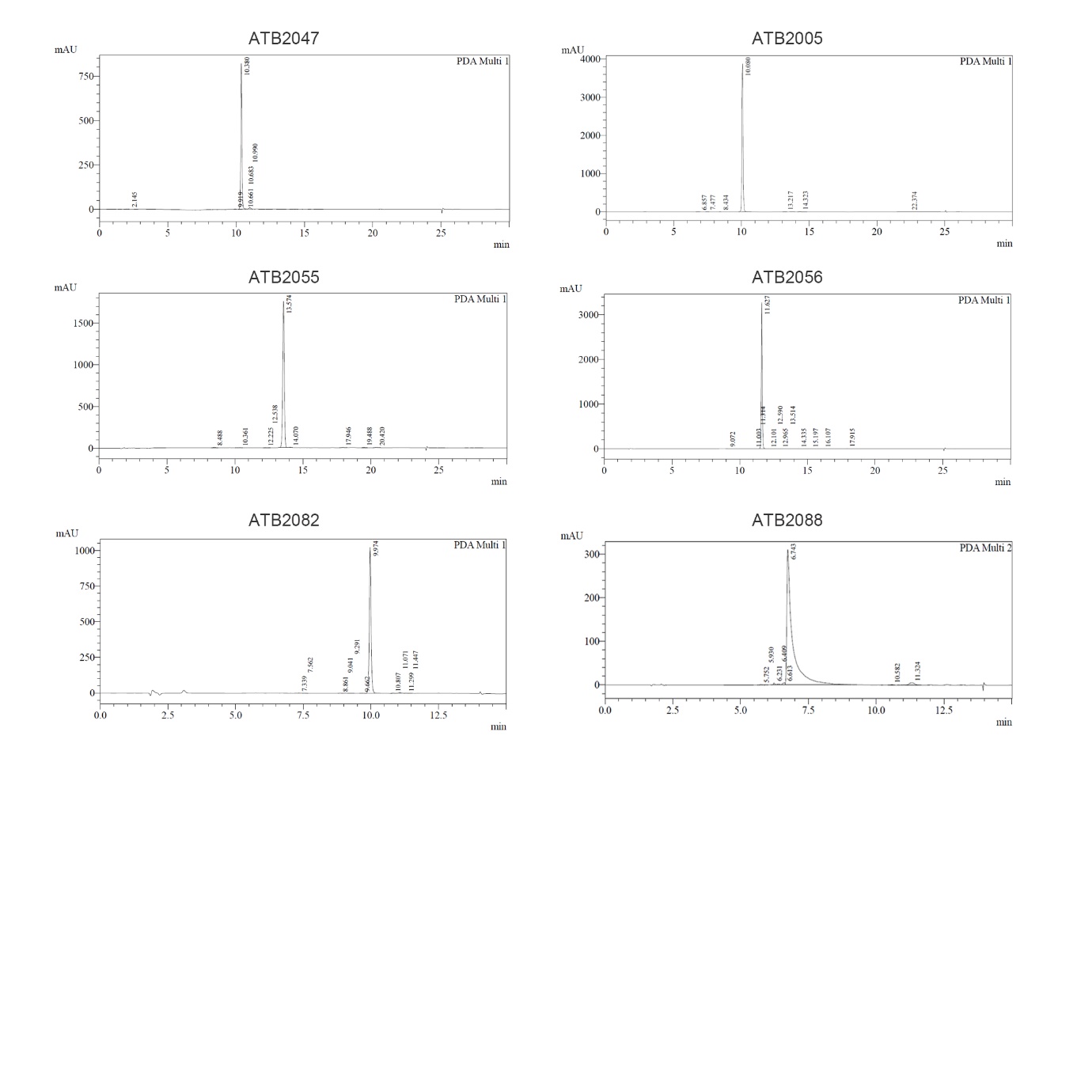

**LC-MS data of AUTOTACS used in this study**

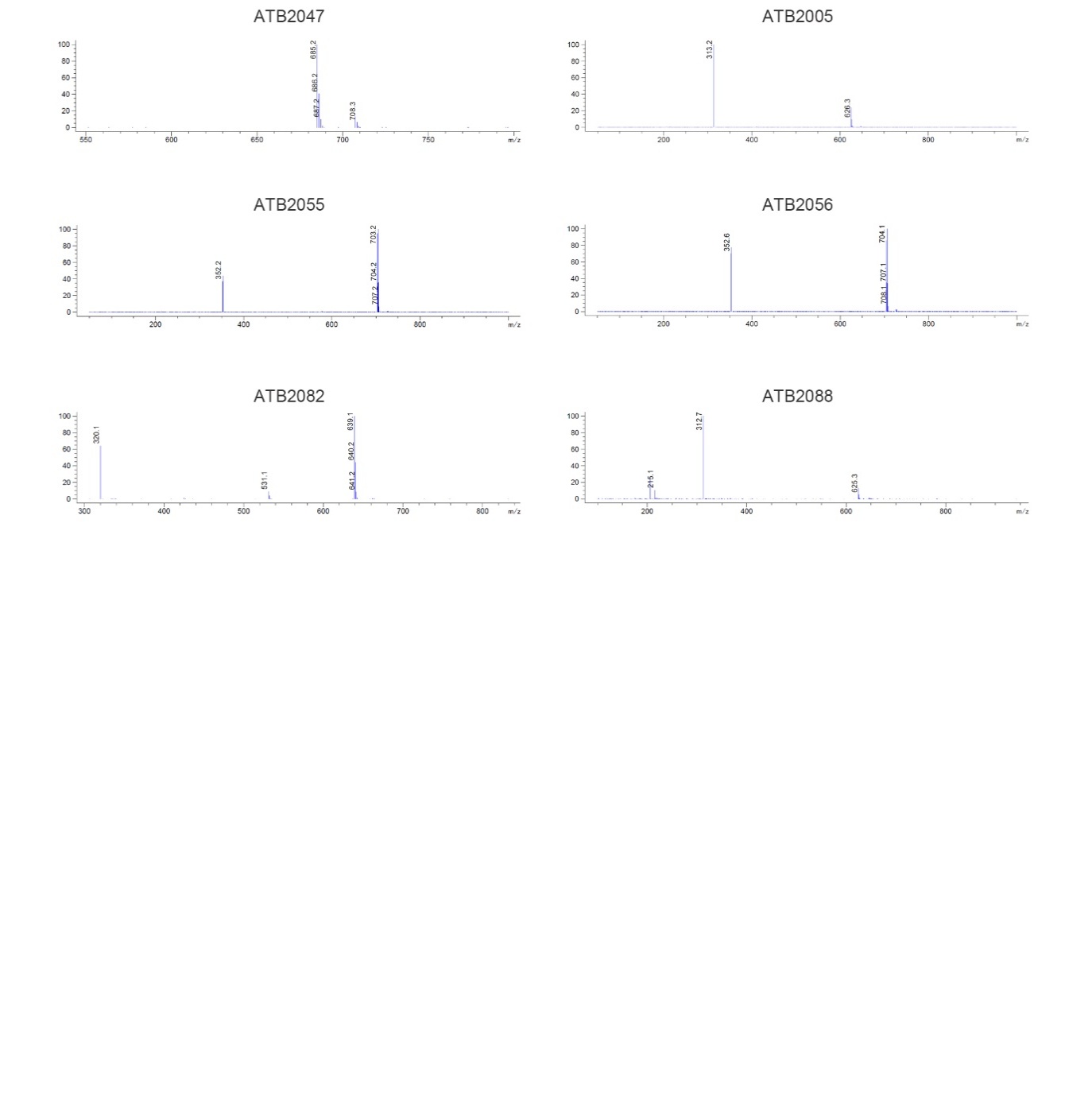
